# Cytostatic hypothermia and its impact on glioblastoma and survival

**DOI:** 10.1101/2021.03.25.436870

**Authors:** Syed Faaiz Enam, Cem Y. Kilic, Jianxi Huang, Brian J. Kang, Reed Chen, Connor S. Tribble, Ekaterina Ilich, Martha I. Betancur, Stephanie J. Blocker, Steven J. Owen, Anne F. Buckley, Johnathan G. Lyon, Ravi V. Bellamkonda

**Affiliations:** Department of Biomedical Engineering, Pratt School of Engineering Duke University; Durham, NC 27705, USA; Department of Radiology, Center for In Vivo Microscopy, Duke University; Durham, NC 27705, USA; Bio-medical Machine Shop, Pratt School of Engineering, Duke University; Durham, NC 27705, USA; Department of Pathology, School of Medicine, Duke University; Durham, NC 27705, USA

## Abstract

Novel therapeutic approaches are needed for patients with glioblastoma (GBM) who otherwise have limited options. Here we studied and deployed non-freezing ‘cytostatic’ hypothermia to stunt GBM growth. This contrasts with ablative, cryogenic hypothermia: a double-edged sword against tumors infiltrating otherwise healthy tissue. We investigated three grades of hypothermia *in vitro* and identified a cytostatic window of 20–25°C. For some glioma lines, 18 h/d of cytostatic hypothermia was sufficient to halt division *in vitro*. Cytostatic hypothermia induced cell cycle arrest, reduced metabolite production and consumption, and reduced inflammatory cytokine synthesis. Next, we fabricated an experimental device to test local cytostatic hypothermia *in vivo* in two rodent models of GBM: utilizing the rat F98 and the human U-87 MG lines. Hypothermia more than doubled the median survival of F98 bearing rats from 3.9 weeks to 9.7 weeks and two rats survived through 12 weeks. All U-87 MG bearing rats that successfully received cytostatic hypothermia survived their study period. Thus, this approach lengthened survival without chemical interventions. Unlike targeted therapeutics that are successful in preclinical models but fail in clinical trials, cytostatic hypothermia affects multiple cellular processes simultaneously. This, alongside reduced cellular division, suggests that opportunities for tumor evolution are reduced and the likelihood of translation to larger species may be more likely. In addition, based on our work, designs, and the literature, engineering a patient-centric device is tangible. Taken together, cytostatic hypothermia could be a novel approach to cancer therapy and eventually serve a valuable role to patients with GBM.

**One Sentence Summary:** Hypothermia influences multiple cellular pathways, can be a safe and effective approach to halt glioblastoma growth, and holds translational promise.

## INTRODUCTION

Despite standard-of-care treatment, patients with glioblastoma (GBM) have a poor median survival of 15–18 months. At best, ~7% survive 5 years after diagnosis *(1, 2)*. This is due to the ineffectiveness of current therapies, resulting in nearly all GBMs recurring. Interestingly more than 80% of recurrences are local *(3–5)*, which provides a role for local therapies. Unfortunately, due to the traumatic effects of treatment, only 20–30% of the recurrences can be resected before recurring once more *(6)*.

The ineffectiveness of current GBM therapies necessitates novel domains of therapeutics, such as manipulating physical phenomena like electric fields, topographical guidance, and temperature. For example, tumor-treating electric fields can disrupt mitosis and have demonstrated a mild effect on overall survival *(7, 8)*. Topological cues of aligned nanofibers can induce directional migration of GBM to a cytotoxic sink *(9)*, and this strategy recently received FDA breakthrough status. In fact, directional GBM migration may also be induced by applying electric fields *(10)*. Both hyper- and hypo-thermia have been used to successfully kill tumor cells *(11, 12)*. Thus, approaches that leverage physics may expand the repertoire of options available to GBM patients.

Hypothermia as a cancer therapy remains relatively under-explored and can be divided into two forms: cryogenic (freezing) and non-cryogenic hypothermia. Currently, our only hypothermia-driven approach against cancer is cryosurgery *(13)*. In the 1940s, neurosurgeon Dr. Temple Fay first applied whole-body hypothermia to limit tumor growth, but this proved to be hazardous and infeasible *(14)*. He then attempted locally freezing tumor in one glioma patient with a cryoprobe tethered to a ‘beer cooler’ *(14)*. However, the degree, duration, intermittency, and follow-up of this case is not detailed, possibly due to hypothermia falling out of favor after it was misused during World War II *(15)*. Subsequently, cryogenic freezing of brain tumors was repeated and cryosurgery was born *(12, 16)*. However, subzero temperatures indiscriminately ablate diseased and healthy tissue *(13, 17)*. Thus, cryogenic freezing offers little advantage over current GBM therapies.

Alternatively, the healthy brain is resilient to non-cryogenic hypothermia *(14, 18, 19)*. Additionally, temperatures of 32–35°C may be neuroprotective after injury *(20)*. This moderate hypothermia reduces cell metabolism, oxygen and glucose consumption, edema, excitotoxicity, and free-radical formation *(20)*. In epilepsy, novel cortical cooling devices can halt seizures in primates *(19, 21)* and intraoperatively in patients *(22)*. Thus, non-cryogenic hypothermia could have therapeutic utility without significant cortical damage.

In this study, we propose non-cryogenic hypothermia to halt brain tumor growth in awake and freely moving rodents. We first identify a window of temperatures that safely halts cell division, and we term this range ‘cytostatic hypothermia’. Our approach validates and expands upon suggestions from *in vitro* studies *(23–28)*, including one from 1959 *(14)*, that hypothermia can reduce cell proliferation. We investigate the effects of the degree and duration of hypothermia on the growth and metabolism of multiple human GBM lines *in vitro*. We also explore the effect of concomitant hypothermia with chemotherapy and chimeric antigen receptor T cell (CAR T) immunotherapy *in vitro*. Next, we computationally model and fabricate an experimental device to deliver cytostatic hypothermia *in vivo* for rats to demonstrate proof-of-concept. We then test the application of cytostatic hypothermia in two GBM rat models. Based on our findings, we propose what a device for patients could look like. Leveraging a fundamental *physical* phenomenon that influences biology (temperature), multiple cellular pathways are modulated simultaneously, and the broad effects may not be species-specific (i.e., they may translate to humans). Based on our evidence and the potential for translation, cytostatic hypothermia could be a much-needed option for patients who would otherwise succumb to GBM.

## RESULTS

### In vitro glioblastoma growth is influenced by hypothermia

To investigate how different degrees of hypothermia affect cell growth rates, we cultured three human GBM cell lines and one rat GBM line at 20, 25, 30 and 37°C. Growth was assessed daily through a custom live-cell imaging and analysis method we developed (Fig. 1A). It uses contrast and edge detection with optimized thresholds to identify tumor coverage as a proxy for growth in a 2D well. All cell lines grew at 37°C (fig. S1A) and became fully confluent within 3– 5 days (Fig. 1B). Reducing the temperature to 30°C reduced growth rates such that a significant difference in cell coverage was observed out to 12 days in three lines. Furthermore, three human GBM lines all showed no growth at 25°C (Fig. 1B, fig. S1B). Interestingly, while F98 (rat GBM) did have significantly reduced coverage at 12 days under 25°C, it required 20°C to halt growth (Fig. 1B, fig. S1C).

**Figure 1:**
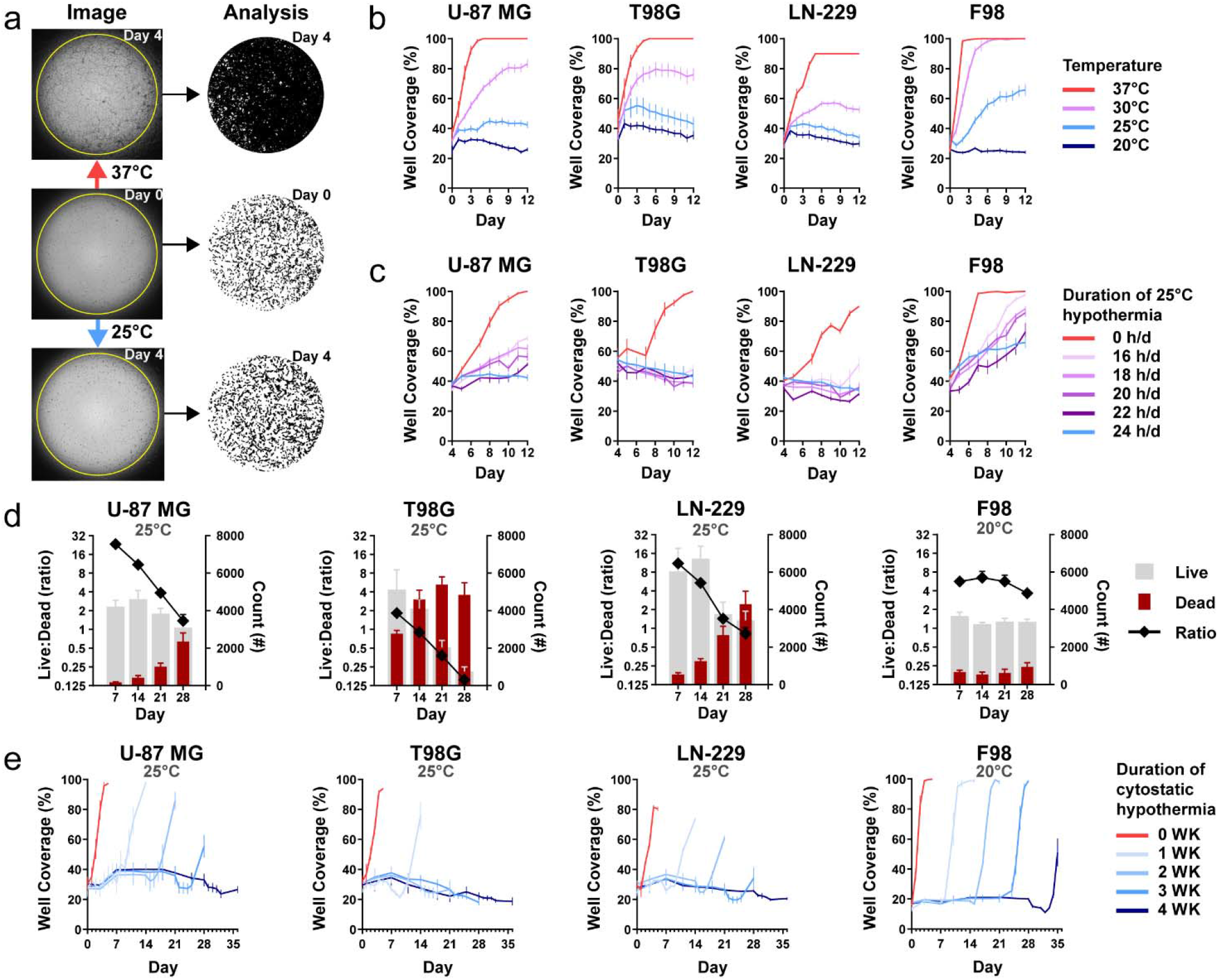
Effect of continuous and intermittent cytostatic hypothermia on GBM cell lines *in vitro*. **(A)** Custom imaging and tumor well coverage analysis assay with photos (left) and resulting masks (right). Cells were incubated overnight in 96-well plates at 37°C (middle panel) and then incubated under either normothermia or hypothermia for 4 days (upper and lower panels). A custom ImageJ macro identified tumor from background and quantified confluence as “Well Coverage (%)”. On the masks, black represents cells and white represents the well. **(B)** GBM growth curves (n = 8) via percent of well coverage at 37, 30, 25, and 20°C; two-way ANOVA with Dunnett’s multiple comparisons test demonstrated *p* < 0.0001 at day 12 between all hypothermia conditions and 37°C across all lines, except for F98 at 30°C (non-significant). **(C)** GBM growth rates under intermittent 25°C hypothermia after 25°C pre-treatment (n = 8). Cells were plated and grown overnight at 37°C, followed by 4 days under 25°C hypothermia (pre-treatment). Beginning day 4, cells were transferred between 37°C and 25°C incubators for varying hours per day (h/d); two-way ANOVA with Dunnett’s multiple comparisons test demonstrated *p* < 0.0001 at day 12 between all durations and 0 h/d across all lines, except for F98 at 16 h/d (*p* = 0.0013). **(D)** Live/Dead assay of GBM lines over time at cytostatic hypothermia temperatures (25°C for U-87 MG, T98G, LN-229 and 20°C for F98) (n = 8). Groups of the same line were stained with fluorescent microscopy-based live/dead dyes. A custom ImageJ macro counted the number of living (grey) and dead cells (maroon) plotted on the right y-axis. A ratio of live:dead was calculated and plotted on the left y-axis. **(E)** Cell viability imaging assay (n = 8). Cells were grown at their cytostatic temperatures for 0, 1, 2, 3, and 4 weeks. After those time points, the cells were moved to 37°C and incubated for 1 more week. Two-way ANOVA with Dunnet’s multiple comparisons test demonstrated significant difference (*p* < 0.0001) at 7 days after 4 WK hypothermia compared to 7 days after 0 WK hypothermia for U-87 MG, T98G, and LN-229 while *p* = 0.0246 for F98; remaining statistics are reported in Supplementary Table 1. All graphs show mean ± standard deviation.

To explore the minimum daily duration of cooling required to maintain cytostasis, we cycled the plates between a 37°C and 25°C incubator for a pre-defined number of hours per day (h/d). The temperature change of the media had a rapid phase followed by a slow phase, reaching the desired temperature within 30–60 minutes (fig. S2A). The growth rates of all cell lines were significantly reduced with as little as 16 h/d of 25°C hypothermia compared to 37°C (fig. S2B). The application of 20 h/d of 20°C hypothermia on F98 also reduced its growth rate compared to 25°C (fig. S2C). To assess whether an ‘induction’ dose of hypothermia made cells more sensitive to intermittent hypothermia, we added a 4-day 25°C pretreatment period. Growth rates were significantly affected (Fig. 1C) such that T98G and LN-229 demonstrated equivalent well coverage on day 12 compared to day 4 after 18 h/d hypothermia (Fig. 1C).

To assess cell viability after prolonged cytostatic hypothermia, we used a Live-Dead assay (Fig. 1D) at predetermined timepoints (7, 14, 21, and 28 d) and, in a separate assay, observed growth after returning the cells to 37°C (Fig. 1E). All GBM lines demonstrated a reduction in the Live:Dead ratio at their cytostatic temperatures (25°C for human lines and 20°C for F98) with T98G being the most affected and F98 being the least affected (Fig. 1D). Similarly, growth lagged after removal of 2 or 3 weeks of cytostatic hypothermia in all lines except for F98. Growth lagged in all lines (including F98) after 4 weeks, with significantly reduced well coverage 7 days after returning to 37°C (Fig. 1E, table S1).

The custom imaging assay also facilitated assessment of morphological features of circularity and average size. Compared to cell circularity after overnight incubation at 37°C (day 0), hypothermia significantly increased U-87 MG and LN-229 circularity (fig. S2D, table S2) while T98G remained unaffected. While F98 circularity significantly decreased with hypothermia over time, it displayed significantly greater circularity under 20°C compared to 25°C. Similarly, average cell size of U-87 MG and T98G were significantly reduced with prolonged hypothermia (fig. S2E, table S3). F98 demonstrated an increase in the average cell size under 25°C, but this was abrogated under its cytostatic temperature of 20°C (fig. S2E, table S3).

### Hypothermia arrests the cell cycle and reduces metabolism and cytokine synthesis

To further investigate the effects of cytostatic hypothermia, we assayed cell cycle, ATP levels, metabolite production/consumption, and cytokine production under hypothermia. Cells grown for 3 days at 37°C were predominantly in the G1 phase of the cell cycle (Fig. 2A). The application of 3 or 7 days of 25°C, induced a significant reduction in both the G1- and S-phases along with an increase in the G2-phase across all cell lines (Fig. 2A). Next, we quantified intracellular ATP to investigate whether ATP production and consumption ceased. Interestingly, intracellular ATP was significantly reduced in T98G but significantly greater in the remaining 3 lines under 25°C hypothermia (Fig. 2B, table S4). Next, we assayed media glucose, lactate, glutamate, and glutamine at 0, 3, 7, and 14 days at 20, 25, 30, and 37°C (Fig. 2C and fig. S3). To account for cell growth under non-cytostatic conditions, the data were normalized to tumor-cell occupied surface area using our imaging assay (Fig. 2C). Results demonstrated that metabolite production and consumption are significantly reduced with any degree of hypothermia. F98 glutamate production per unit cell area was not significantly different at day 3 (Fig. 2C) but remained low under 20°C (fig. S3D). Similarly, expression of inflammatory cytokines IL-6 and IL-8 from human GBM lines were significantly lower after 6 or 10 days of hypothermia (Fig. 2D). Interestingly, a mild but significant increase of both cytokines was observed after the first 3 days of 25°C from T98G, but it subsided by day 6. We did not detect expression of anti-inflammatory cytokines IL-4, IL-10, or the inflammatory cytokine IFNγ under any condition.

**Figure 2:**
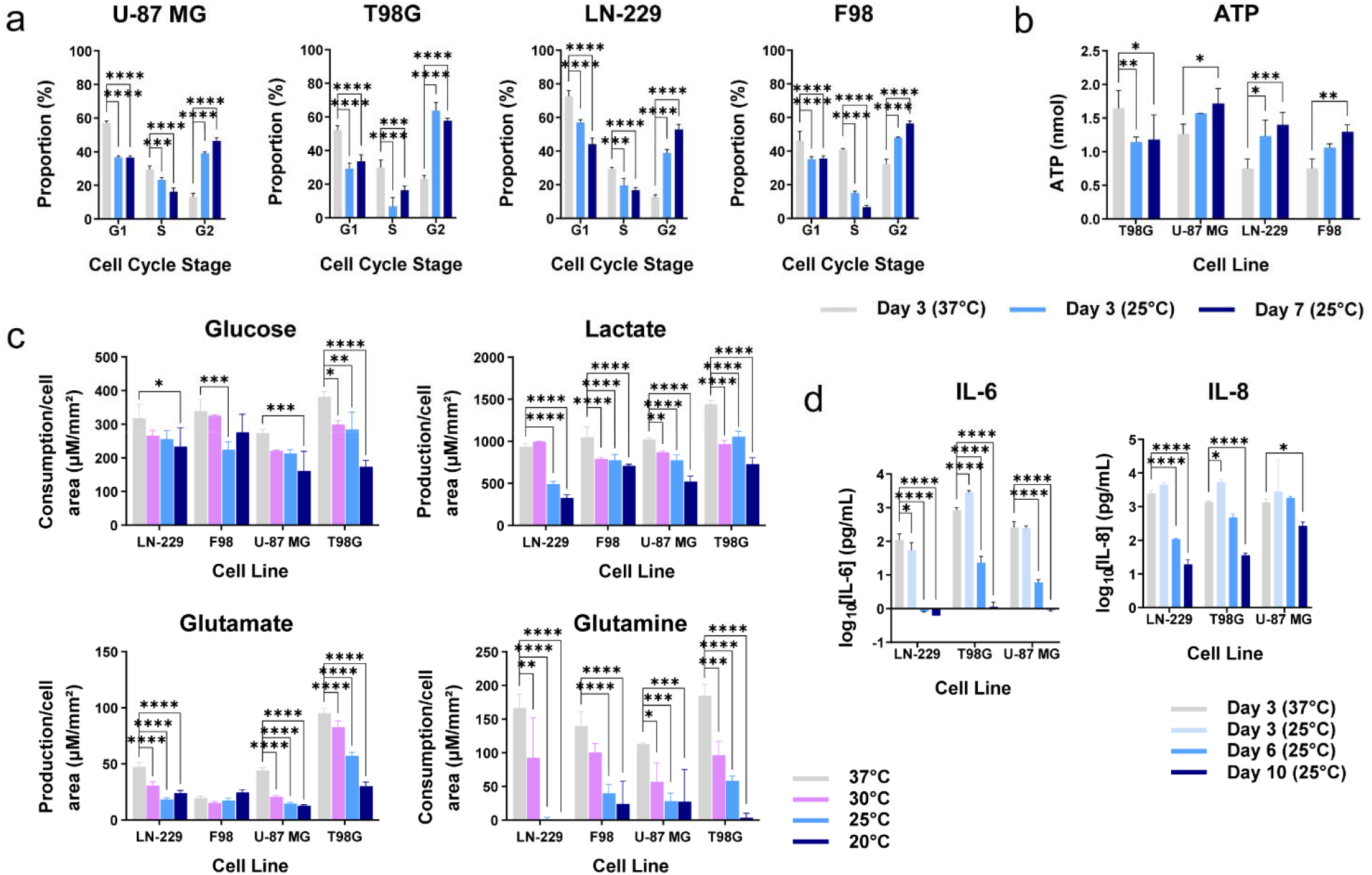
Effects of hypothermia on tumor cell cycle, metabolism, and cytokine synthesis. **(A)** Percentage of cells in each stage of the cell cycle after either 3 days of 37°C, 3 days of 25°C, or 7 days of 25°C (n = 3). **(B)** Amount of intracellular ATP from the tumor cell lines when grown for 3 days at 37°C, 3 days at 25°C, or 7 days at 25°C (n = 3). Specific adjusted *p*-values are provided in Supplementary Table 4. **(C)** Metabolite production or consumption per cell surface area across cell lines and temperatures after either 3 days of 37°C or 3 days of hypothermia (30°C, 25°C, 20°C) (n = 3). **(D)** Concentration of cytokines (IL-6 and IL-8) collected in the media at different time points and temperatures (n = 3). All studies used two-way ANOVAs with post-hoc Dunnett’s multiple comparison test (\**p*<0.05, \*\**p*<0.01, \*\*\**p*<0.001, \*\*\*\**p*<0.0001). All graphs show mean ± standard deviation.

### Adjuvants can function with cytostatic hypothermia *in vitro*

We next assessed whether cytostatic hypothermia would dampen temozolomide (TMZ) chemotherapy or CAR T immunotherapy *in vitro*. Human GBM lines received TMZ at one of three concentrations (0, 500, or 1000 μM) and at either 37°C or 25°C followed by media replacement and incubation at 37°C (for details, see Methods). All three lines had reduced well coverage 12 days after completion of treatment with 1000 μM of TMZ compared to no TMZ (DMSO only) under 25°C (fig. S4A). Additionally, both TMZ and hypothermia halted cells in the G2-phase of the cell cycle (fig. S4B). U-87 MG growth was significantly reduced on day 13 by combined TMZ and hypothermia compared to TMZ alone (at 37°C) (Fig. 3A). Growth of T98G, a TMZ-resistant line, was significantly reduced by hypothermia alone and unaffected by 500 and 1000 μM of TMZ alone (at 37°C) (Fig. 3B). However, the combination of 500 or 1000 μM TMZ with 25°C hypothermia had significantly delayed growth by day 13. Similarly, LN-229 growth was significantly reduced by both TMZ and hypothermia individually, and the combination resulted in an enhanced effect (Fig. 3C).

**Figure 3:**
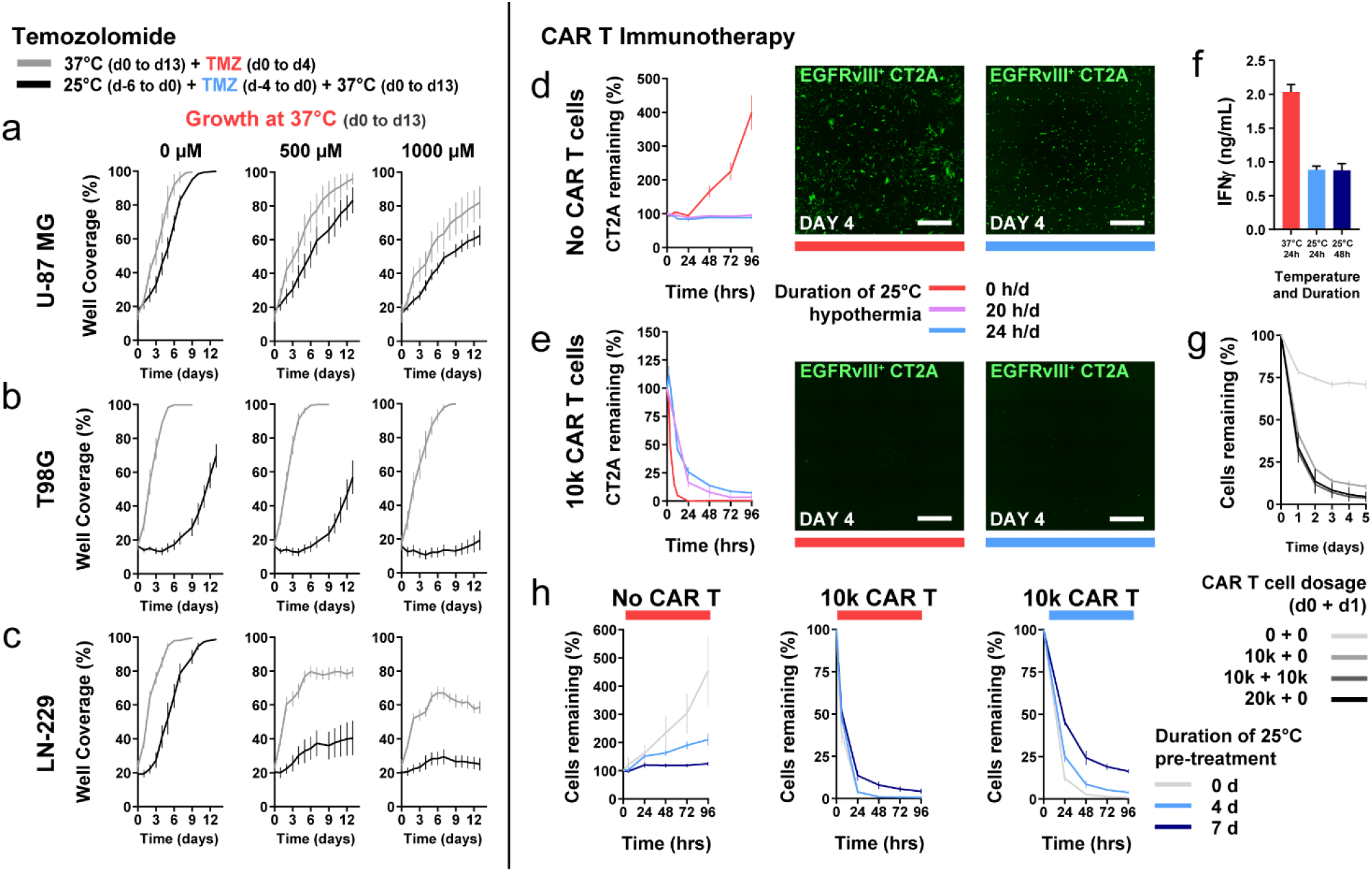
Use of cytostatic hypothermia with chemotherapy and CAR T immunotherapy. **(A–C)** Growth curves of U-87 MG, T98G, and LN-229 at 37°C with or without TMZ (DMSO only or 500 and 1000 μM TMZ) and either after or without hypothermia. Plotted gray line represents growth with concomitant TMZ chemotherapy from days 0–4 at 37°C and subsequent growth in fresh media. Plotted black line represents growth at 37°C in fresh media *after* completion of TMZ and 25°C hypothermia treatment (n = 8). Two-way ANOVA with Sidak’s post-hoc test demonstrated significant differences between the 37°C and 25°C groups on day 13 in **(A)** U-87 MG at 500 μM (*p* = 0.0207) and 1000 μM (*p* = 0.0292), **(B)** T98G at 500 μM (*p* < 0.0001), and 1000 μM (*p* < 0.0001), and **(C)** LN-229 at 500 μM (*p* < 0.0001), and 1000 μM (*p* < 0.0001). There were no significant differences on day 0 between 37°C and 25°C for each cell line. **(D–E)** GFP^+^ EGFRVIII^+^ CT2A tumor cells remaining after treatment without **(D)** and with 10,000 CAR T cells **(E)** under normothermia, or continuous or intermittent hypothermia (20 h/d) (n = 4). Representative images of remaining tumor cells at day 4 after beginning treatment. White scale bars = 1000 μm. **(F)** IFNγ quantification from media with EGFRvIII^+^ CT2A cells and CAR T cells. One-way ANOVA with Dunnett’s post-hoc test demonstrated a significant difference (*p* < 0.0001) between the 37°C and each of the 25°C groups. **(G)** Remaining CT2A cells with higher or multiple CAR T cell dosage over 2 days under 25°C hypothermia. **(H)** Remaining CT2A cells after pretreatment with 25°C hypothermia for 0, 4, or 7 days, followed by addition of 0 or 10,000 CAR T cells, and plate moved to either 37°C (left and middle) or 25°C (right). All graphs show mean ± standard deviation.

As CAR T immunotherapies against glioblastoma are under investigation *(29)*, we assessed whether CAR T induced cytotoxicity (of EGFRvIII^+^ CT2A mouse glioma cells) was possible under cytostatic hypothermia. In the absence of CAR T cells, CT2A cell growth was inhibited by both 24 and 20 h/d of 25°C hypothermia (Fig. 3D). Expression of the chimeric antigen receptor on T cells enabled killing of GFP^+^EGFRvIII^+^ CT2A cells and not GFP^+^EGFRvII^-^cells at 37°C (fig. S4C). With the addition of 10,000 CAR T cells (5:1 ratio of T cell:tumor), EGFRvIII^+^ cells were eradicated within 24 hours at 37°C (Fig. 3E). Under 25°C hypothermia, 10,000 CAR T cells significantly reduced the glioma cells to 25% of the original count within 24 hours and to <10% by 96 hours. Reducing the hypothermia dosage to 20 h/d further reduced the remaining tumor percentage (Fig. 3E). Reduced IFNγ was detected in the media when immune mediated killing took place under hypothermia (Fig. 3F). Under hypothermia, adding a second dose of CAR T cells after 24 hours brought the remaining tumor to <5% of the original cell count (Fig. 3G). Pretreating CT2A cells with 25°C hypothermia for 4–7 days reduced their subsequent growth rate at 37°C (Fig. 3H, left). However, this reduced their susceptibility to CAR T cells at 37°C (Fig. 3H, middle) and 25°C (Fig. 3H, right).

While these results suggest that cytostatic hypothermia may function with adjuvants *in vitro*, its potential *in vivo* requires optimization and characterization, detailed in the Discussion. Initial clinical trials would evaluate cytostatic hypothermia efficacy alone in recurrent and resistant GBMs (as a last resort). Thus the *in vivo* potential for *concomitant* hypothermia could be postponed as the field matures and a patient-centric device is established.

### Hypothermia can be delivered *in vivo* and reduces GBM tumor growth

Given that hypothermia of 20–25°C halted cell division *in vitro* (Fig. 1a), we conceived a tool to deliver hypothermia *in vivo* to demonstrate proof-of-concept for cytostatic hypothermia. Using the finite-element method, we computationally modeled local intracranial hypothermia using parameters from prior studies (table S5) *(30–33)*, Pennes’ bio-heat equations, temperature-dependent blood perfusion, and a previously modeled rat brain *(30)* (Fig. 4A and fig. S5A). A single 1-mm wide gold probe was modeled invaginating into the brain with the goal of cooling a spherical volume of tissue (representing either a bulk tumor or the extent of an infiltrating tumor) via a Peltier plate (Fig. 4B and fig. S5B). The simulation demonstrated that a single probe creates a temperature gradient in the adjacent tissue (Fig. 4B). With the probe brought to ~12°C, the periphery of a 1-mm radius region reached 25°C (Fig. 4B) and reached >35°C within 4 mm (Fig. 4C, left). Cooling was greater across the x-axis (of a coronal plane) than down the z-axis (Fig. 4C, left). The extent of perfusion (ranging from 0.73 to 3.96 times brain perfusion) had a small effect on cooling even a larger 1.5-mm radius spherical region (Fig. 4C, middle). To achieve hypothermia in a 1.5-mm radius region, 125mW of heat pulled was sufficient to attain 25°C at the periphery (Fig. 4C, right). Pulling 150mW brought the probe to below 0°C, but the tissue was at 15°C within 0.5 mm from the probe (Fig 4C, right). A time-dependent study using the models suggested that pulling 100-125mW can bring the maximum tumor temperature to its target within 1–2 minutes (fig. S5C).

**Figure 4:**
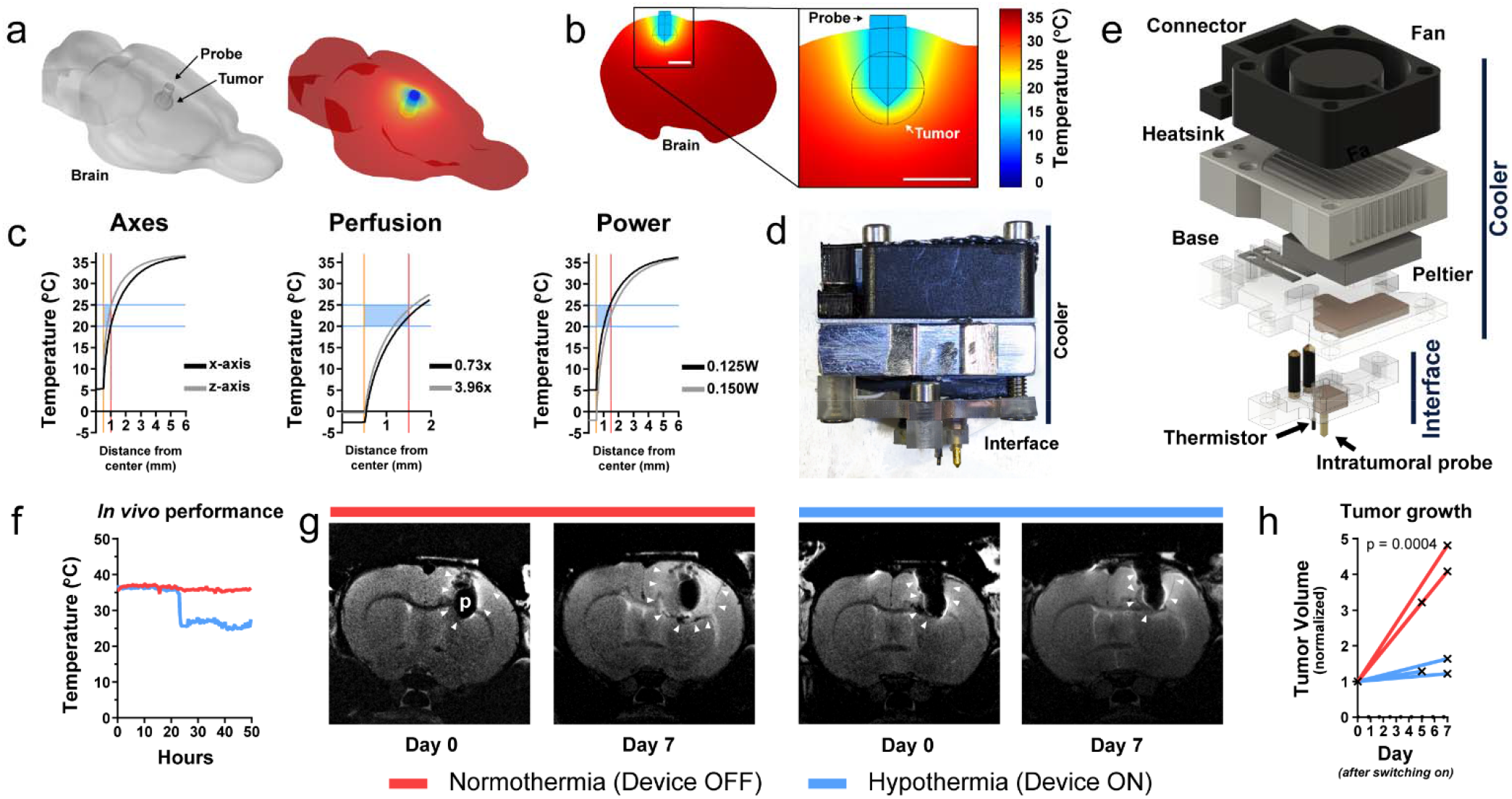
Finite-element analysis and device testing for cytostatic hypothermia delivery. **(A)** Left: 3D model of rat brain with a probe and subcortical spherical tumor. Right: finite-element simulation of temperature on the brain surface. **(B)** Slice and magnified inset from finite-element model of rat brain with local hypothermia. A 1-mm width probe in a 1-mm radius tumor cooled to 12°C enabled tumor periphery to reach 25°C. **(C)** Finite-element analysis of hypothermia varying parameters. Center of tumor and probe lie at x=0. Gold bar indicates surface of gold cooling probe. Red bar indicates surface of tumor. Blue bars and box show cytostatic range of temperature. Initial brain perfusion was set at 0.019333 s^-1^. Left: Comparison of extent of cooling of a 1-mm radius tumor from nearest probe surface in the x- (black) vs. z-axis (gray) of a coronal plane. Middle: Comparison of varying tumor (1.5-mm radius) perfusion relative to brain perfusion (0.73x, black to 3.96x, grey). Right: Varying heat energy withdrawal on cooling a 1.5-mm radius tumor. **(D)** Thermoelectric cooling device consisting of a lower Interface for tissue interface and an upper, removable Cooler for air-based heat exchange. **(E)** Exploded 3D render of thermoelectric device. Cooler consists of a fan, heatsink with cover, Peltier plate, and a base with a copper plate for thermal contact to Peltier and Interface (with thermal paste), and steel shims for thermistor contact. Interface consists of copper plate for thermal contact to Cooler, gold cooling probe as tissue interface, and thermistor with wires around insulated brass screws. Cooler is attached to the Interface by screws. **(F)** *In vivo* temperature measurement from the thermistor of the Interface, 1.5 mm from the probe. Here, both rats possessed a device, but only one was switched on (blue). **(G)** Representative T2-weighted MR images of F98 tumor growth on Day 0 (just prior to switching the device on) and Day 7 in the brains of Fischer rats. The red bar indicates the device was off (normothermia) between images, while the blue bar indicates the device was on (hypothermia). White arrows indicate tumor boundaries; ‘p’ = probe. **(H)** F98 tumor volume measured from MRI and normalized to Day 0 volume (n = 3). Each line represents one rat. Mixed-effects analysis was conducted to compare the groups.

Next, to test cooling *in vivo*, we constructed an experimental device that consists of a removable ‘Cooler’ and an implantable ‘Interface’ (Fig. 4D and fig. S6). The Cooler uses an electrically powered thermoelectric (Peltier) plate secured with thermal paste between a copper plate (embedded in a polycarbonate base) and an aluminum heatsink (Fig. 4E). A fan, attached above, enhances convective heat transfer. The Interface attaches to the skull, has an intratumoral gold needle soldered to a second copper element, and a thermistor wired to insulated brass screws (fig. S6, H and I). The copper parts make contact to transfer heat from the Interface to the Cooler (fig. S6L). This Cooler is powered through an external power supply. Temperature is measured by the thermistor connected to a voltage-divider circuit, Arduino, and a computer (fig. S7).

All rats were inoculated with F98 tumor cells, and tumor-take was confirmed via T2-weighted MRI 1 week after. The following day, the Interface was implanted (demonstrated in fig. S8, A and B) eventually followed by attachment of the Cooler. A custom cage setup was developed to enable free movement of the rats (fig. S8, C to E) *(31)*. Once switched on, the device was able to bring tissue 1.5 mm from the surface of the probe to 25°C (Fig. 4f). Devices did not consistently reach 20°C due to inefficiencies of the heat-exchange mechanism. Interfaces embedded with an MRI-compatible thermistor enabled F98 tumor growth assessment via MRI under normothermia (device off) and hypothermia (device on) (Fig. 4G). MRI image analysis revealed a significant difference in tumor volume growth at 1 week under hypothermia versus tumors at normothermia (Fig. 4H). Due to high failure rates of the MRI-compatible thermistor, subsequent studies used an MRI-incompatible thermistor with tighter tolerance and greater reliability. These preliminary studies demonstrated that *in vivo* local cytostatic hypothermia was both possible and plausibly effective.

### Cytostatic hypothermia extends survival of rodents with GBM in two models

As local hypothermia reduced *in vivo* tumor growth rate in preliminary imaging studies, we investigated its effect on animal survival. In a model with the aggressive F98 line in Fischer rats, we investigated whether survival would increase despite being unable to reach cytostatic temperature for F98 (20°C). After confirmation of tumor-take with GFP^+^ F98 cells (fig. S9A), Interface implantation (example in fig. S8b), and Cooler attachment, rats were randomly assigned to two groups. One group (n = 9) had their devices switched on to deliver hypothermia while the other groups’ (n = 9) remained off. Of the rats receiving hypothermia, only 2 reached near or below 20°C (1.5 mm from the probe, fig. S9B), the cytostatic temperature observed for F98 cells. All rats not receiving hypothermia required euthanasia with a median time of 3.9 weeks (Fig. 5A). After a short phase of recovery from surgery, rats receiving hypothermia did not exhibit obvious signs of weakness or distress, maintained an appetite, and gained weight (Movies 1 and 2, and fig. S9c). In 3 out of 450+ occasions of switching the devices on (after weighing and maintenance of rats), a seizure was observed that abated by turning off the device and did not recur when hypothermia was gradually resumed over a few minutes. With successful application of hypothermia, the median survival for 6 (out of 9) rats was extended to 9.7 weeks (Fig. 5A). The Interface of one rat dislodged prematurely at week 7 and the point was censored. The remaining 2 rats survived the full 12-week study period; these were also the ones in which *cytostatic* hypothermia was achieved (fig. S9B). Thus, the application of hypothermia significantly increased survival (*p* < 0.0001) compared to the normothermia group.

**Figure 5:**
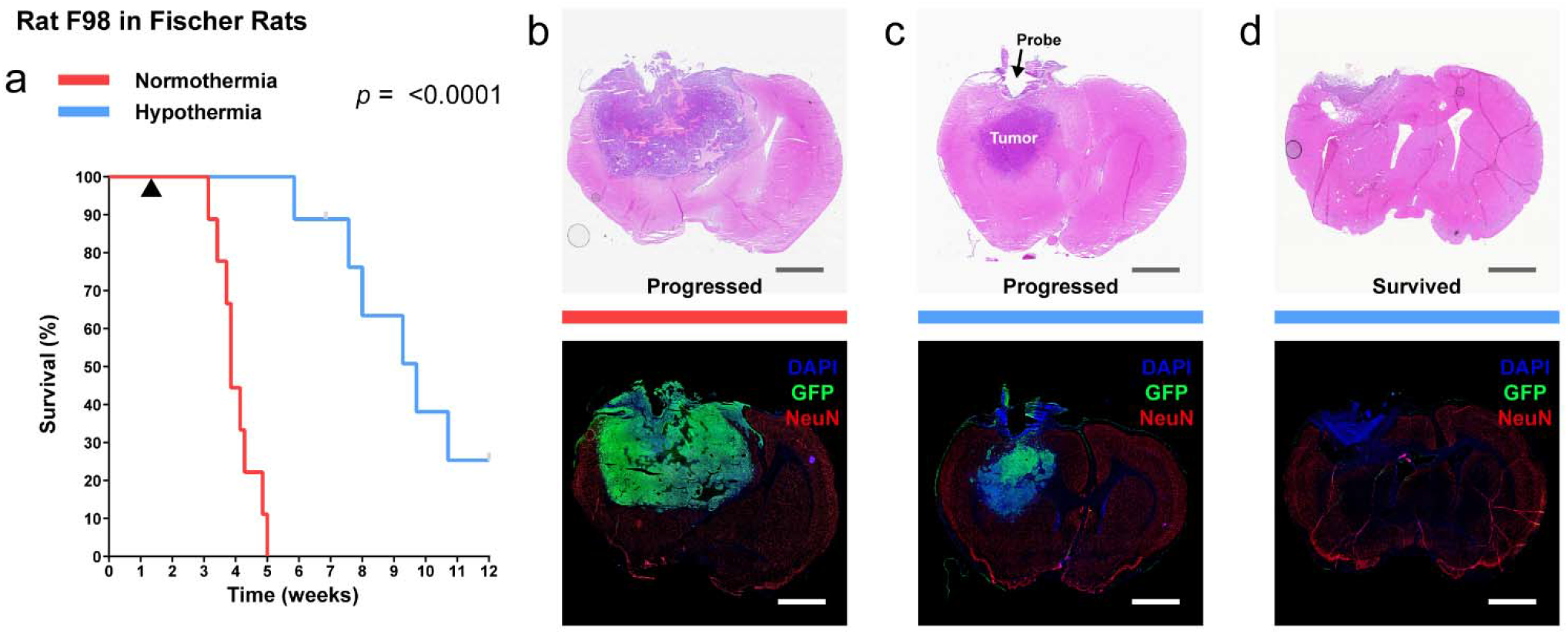
Delivery of local cytostatic hypothermia in Fischer rats inoculated with F98 GBM. **(A)** Kaplan-Meier survival plot of rats with the device switched off (red, n = 9) or switched on to deliver hypothermia (blue, n = 9). Gray bars indicate censored rats. Black triangle indicates the start of hypothermia treatment. Groups were compared with the log-rank Mantel-Cox test. Median survival was calculated for 9 rats in the normothermia group and 6 rats in the hypothermia group. Statistics are for all rats. **(B–D)** Coronal brain sections stained with hematoxylin and eosin (H&E, top) and immunohistochemical markers, DAPI, GFP, and NeuN, (IHC, bottom) to show extent of tumor. Gray and white scale bars = 2.5 mm. **(B)** Representative section from a rat in the normothermia group. **(C)** Representative from a rat in the hypothermia groups in which tumor progressed and rat reached euthanasia criteria. Tumor visible in the z-direction from probe **(D)** Representative section from a rat in the hypothermia group that survived through the study period.

Histology of rats receiving normothermia demonstrated large tumors with leptomeningeal infiltration, tumor-intrinsic necrosis and apoptosis, high Ki-67 activity, and peri-tumoral and ipsilateral GFAP^+^ gliosis (Fig. 5B, fig. S10, Supplementary Tables 6 and 7). With hypothermia, all rats showed necrosis immediately around the probe, with associated inflammation consisting of CD45^+^ mononuclear inflammatory cells and neutrophils (fig. S10F, table S6). Non-neoplastic brain tissue, at the border of the lesions, showed no neuronal loss or parenchymal destruction: NeuN and GFAP expression were always intact (fig. S10A to D, table S6). GFAP demonstrated reactive perilesional gliosis with varied extents of ipsilateral gliosis, and normal patterns of glial expression elsewhere in the brain. Thus, there was no evident difference in the extent of perilesional gliosis in normothermic and hypothermic conditions. The six rats receiving 25°C hypothermia had tumors adjacent to the region of hypothermia (Fig. 5C, table S6). The two rats receiving 20°C (survivors) did not demonstrate a mass (Fig. 5d). They did have a few nonproliferative (Ki-67^-^) GFP^+^ cells (fig. S9D and E) or a rim of viable tumor (table S6).

Next, we assessed cytostatic hypothermia on human U-87 MG inoculated in RNU rats. After confirmation of tumor-take (fig. S11A), Interface implantation (fig. S8B), and Cooler attachment, 6 rats had their device switched on and 5 were kept off (rats were randomly assigned to each group). The devices of 4 treatment rats reliably delivered 25°C hypothermia (fig. S11B), the cytostatic temperature observed for U-87 MG. The thermistors of two rats failed early and treatment of one may not have been reliably delivered (“N06”). All rats receiving normothermia required euthanasia with a median time of 3 weeks (Fig. 6A). As previously, all rats undergoing treatment did not demonstrate signs of distress while receiving hypothermia, had a strong appetite, and gained weight (Movies 1 and 2, fig. S11C). RNU rats were more social and active than Fischer rats and this persisted despite hypothermia. Because of this, their devices and cables required frequent maintenance and replacement with intermittent failures. Eventually, Interface implants started to break off by the 7^th^ week, points were censored, and the study was terminated at 9 weeks (Fig. 6A). We attempted reimplantation in two rats (including “N06”), but this resulted in complications and days without treatment and the rats were censored. None of the rats in the hypothermia arm met euthanasia criteria within the study period (Fig. 6A) demonstrating a significant increase in survival (*p* = 0.0007). Histology demonstrated that rats not receiving hypothermia had large tumors with at least some tumor-intrinsic necrosis (Fig. 6B, tables S6 and S7). The hypothermia rats again demonstrated necrosis with inflammatory cells immediately around the probe and intact NeuN and GFAP staining in the non-neoplastic tissue at the interface of the lesion with brain parenchyma (tables S6 and S7). In the rats receiving re-implantation (and absence of treatment for a few days), a tumor mass and the presence of blood were visible (Fig. 6C, table S7). The remaining four hypothermia rats did not exhibit any definitive tumor away from the probe region (Fig. 6D, table S6). Together, these *in vivo* F98 and U-87 MG studies suggest that delivery of hypothermia, and especially *cytostatic* hypothermia, can prolong survival.

**Figure 6:**
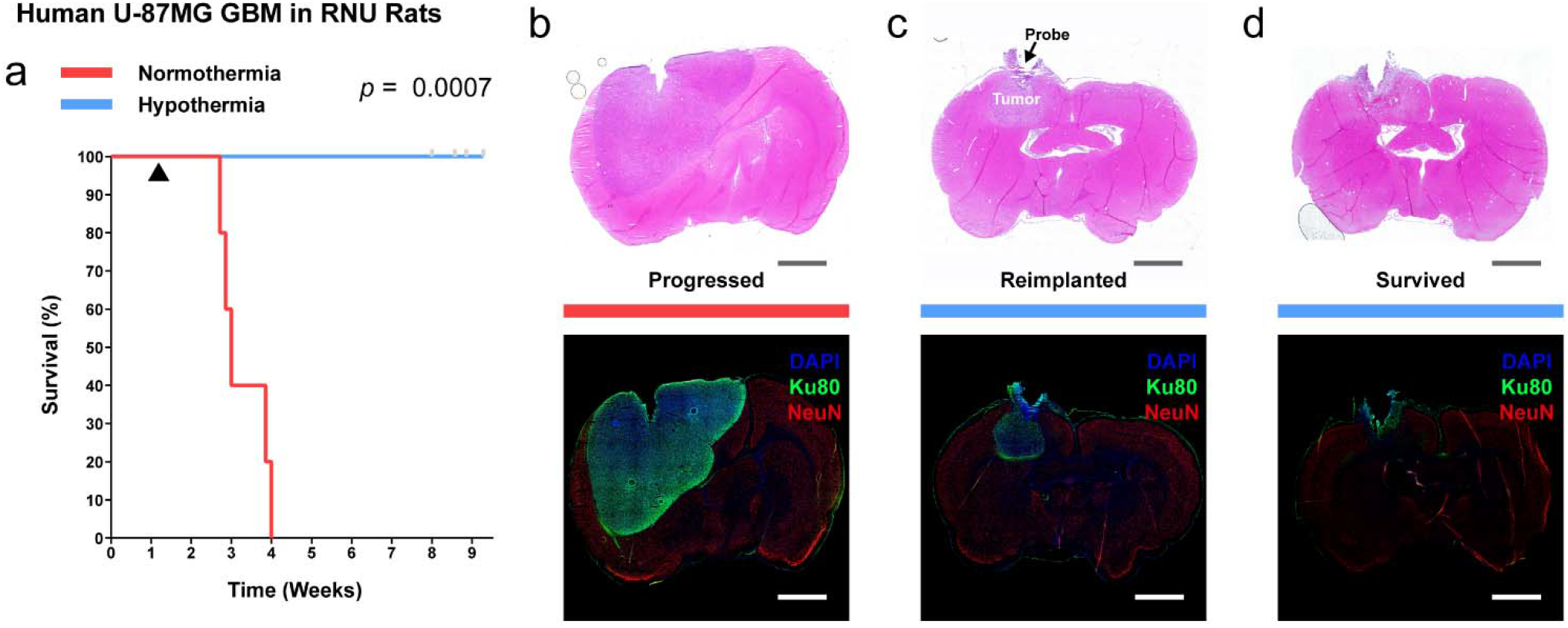
Delivery of local cytostatic hypothermia in RNU rats inoculated with human U-87 MG. **(A)** Kaplan-Meier survival plot of rats with the device switched off (red, n = 5) or switched on to deliver hypothermia (blue, n = 6). Gray bars indicate censored rats. Black triangle indicates start of hypothermia. Groups were compared with the log-rank Mantel-Cox test. **(B–D)** Coronal brain sections stained with H&E (top) and IHC (bottom) to show extent of tumor. Gray and white scale bars = 2.5 mm. **(B)** Representative sections from a rat in the normothermia group. **(C)** Representative sections from a rat in the hypothermia groups that had poor treatment management or multiple device failures and grew a tumor in the z-axis (n = 2). **(D)** Representative sections from a rat in the hypothermia group that did not grow a separate tumor mass (n = 4) and survived the study period.

## DISCUSSION

These studies demonstrate the potential of hypothermia for preventing tumor growth. Through regimented *in vitro* studies, we defined a window of growth-halting temperatures, assessed their effects on cells, and explored strategic intermittent timelines to improve the logistics of implementation. *In vitro* studies also indicated that adjuvants may function with hypothermia but require extensive characterization for *in vivo* applicability. We computationally modeled hypothermia delivery and fabricated experimental devices to administer and monitor intracranial hypothermia, including one which was MR imaging compatible. In two rodent models of GBM, hypothermia extended survival while letting the animals behave normally and eat and move freely (Fig. 5 and Fig. 6, and Movies 1 and 2). We suggest that the range of safe but growth-halting temperatures we term ‘cytostatic hypothermia’ carries translational advantages and could one day be an option for patients.

We established cytostatic temperatures ranging from 20–25°C (Fig. 1B and Fig. 6D) for 5 glioma cell lines and observed multiple biochemical changes. This expands upon and validates findings suggesting that mild hypothermia can reduce cell division *(14, 23–27)*. This evidence is substantial, but future studies with patient-derived lines can investigate more precise cytostatic temperatures and explore mechanistic differences. In addition, we observed that intermittent hypothermia, especially after an ‘induction’ dose, could be equally effective (Fig 1c). Future studies could evaluate the mechanism of induction, but it may be a matter of cells arresting in the G2-phase (Fig. 2A) and intermittent normothermia not providing enough energy or time to complete division. With prolonged hypothermia, and cell cycle arrest, a decrease in viability was apparent (Fig. 1, D and E). This may not be ideal as healthy cells, although more resilient to hypothermia *(24)*, may die too. Potential mechanisms include Na^+^ or Ca^2+^ accumulation or increased aquaporin-4 expression *(32–34)* with cell swelling. To circumvent this, intermittent hypothermia may be equally cytostatic (Fig. 1C) but potentially less cytotoxic. Intriguingly, we discovered that F98 cells required 20°C hypothermia to halt division (Fig. 1B) and this was witnessed in its morphology (fig. S2, D and E) while it retained viability (Fig. 1D). A mechanistic cause was not elucidated but deeper metabolomic or transcriptomic analyses *(35)* could help. Our brief examination of cytokine and metabolite production and consumption demonstrated reduced activity at all degrees of hypothermia. However, as in a study of hypothermia for organ transplant preservation, accumulating ATP in most lines (Fig. 2b) also suggests that one process (e.g., consumption) can be more affected than the other (e.g., production). Cytostatic hypothermia could nevertheless be an alternate approach to pharmaceutical and dietary efforts to starve tumors of resources by reducing glucose consumption and lactate production *(36–38)*. Similarly as glutamate is known to have a role in glioma growth *(39)*, and inflammatory cytokines IL-6 and IL-8 (Fig. 2C) are strongly associated with glioblastoma invasiveness *(40)*, reducing their production might be beneficial. Thus, evidentially, hypothermia can be cytostatic and has broad effects on multiple cellular pathways simultaneously.

Our preliminary examination of hypothermia with TMZ chemotherapy (Fig. 6, A to C) and CAR T immunotherapy *in vitro* suggested that concomitant hypothermia may be a future strategy. Literature suggests that hypothermia can facilitate radiotherapy as well *(41)*. We observed that TMZ and cytostatic hypothermia together reduced growth of all three cell lines tested. This includes T98G cells which are typically TMZ resistant due to the enzyme MGMT *(42)*. This synergy needs to be studied but could be due to increased hydrolysis of TMZ to its active ion, reduced MGMT expression, or compromising effects of hypothermia. On the immunotherapy front, as anticipated from whole-body hypothermia studies *(43)*, hypothermia reduced IFNγ production (Fig. 6F) and immune-mediated killing (Fig. 6E). Nevertheless, albeit over 4 days vs 1 day, CAR T cells still eradicated most glioma cells under cytostatic hypothermia (Fig. 6E). Intermittent hypothermia improved this efficacy (Fig. 6E) suggesting future paradigms of applying cytostatic hypothermia in between CAR T treatments under normothermia. However, there are numerous caveats and limitations that remain to be explored. The three human lines all had different responses to TMZ + hypothermia. For *in vivo* intervention, the effect of hypothermia on vasoconstriction and intravenous drug or cell delivery needs to be studied. Local delivery could be attempted but necessitates designing the device to enable injections. Critically, however, initial clinical trials will likely be conducted only on recurrent and resistant GBMs (*after* standard-of-care) and using only cytostatic hypothermia as a last resort. Thus the *in vivo* potential for *concomitant* hypothermia would be relevant once the field matures, therapeutic and biological mechanisms are better understood, and a patient-centric device is established.

Fortunately, the literature and our studies present ample evidence to suggest that cytostatic hypothermia alone may be safe in the brain. Intracerebral cryoprobes and cortical cooling devices in primates and patients have been safe to use *(14, 16, 19, 21, 22)*. In fact, as we found intact neurons and glia at 20-25°C, another study locally cooling feline cortex down to 3°C, 2 h/d, for 10 months demonstrated neuronal preservation and minimal histological changes *(18)*. Cooling the feline visual cortex intermittently for >3 years also demonstrated this and found visual evoked potentials were fully intact above 24°C and absent only below 20°C *(19)*. Fortuitously, our data demonstrated all GBM lines halted at 20°C or above. Computational modelling also demonstrated that brain perfusion kept cooling diffuse but relatively local to the region of interest (Fig. 3c), which is also seen in the literature *(44)*. A minor reduction in neuronal activity could also have therapeutic advantages too given the recent discovery that inhibiting neuro-glial electrical conduction reduces glioma growth *(45, 46)*. On the behavioral level, we observed that rats generally performed ‘normally’: they responded to stimuli, ate food, gained weight, and moved freely (Movies 1 and 2). Additionally, while hypothermia treatment exhibited tumor necrosis with leukocytic inflammation in the immediate region of the probe, there was no evident parenchymal damage to the adjacent brain. Even at the interface with the lesion, no neuronal or glial loss was detected on immunostaining for NeuN and GFAP respectively (fig. S10, A to D, and table S6). This provides *in vivo* evidence to an *in vitro* study suggesting that hypothermia can selectively protect non-neoplastic cells *(24)*. There was no evidence of infarction, infection, or herniation and reactive gliosis was limited to the ipsilateral hemisphere with no difference between treatment and controls (table S7). However, adverse effects of hypothermia are still possible from reduced neural firing and cellular swelling; these may be resolved by gradual temperature adjustments, intermittent hypothermia, administering molecular inhibitors *(32–34)*, and plausibly the use of adjuvants to reduce the total length of therapy. The extent of safety and behavioral analysis with long-term treatment applied to functional cortex remains to be studied. Such studies could investigate subtle abnormalities and assesses any plasticity-induced recovery over time *(47)*. Thus, while remembering the adverse effects of surgery, chemotherapy, and radiation, cytostatic hypothermia could be relatively benign as is already demonstrated in the literature and through our *in vivo* studies.

Perhaps the most exciting finding from this work was that a physical phenomenon extended survival of rodents bearing GBM. In these proof-of-concept studies, hypothermia was delivered to bring the periphery of a bulk tumor (U-87 MG) and one with leptomeningeal infiltration *(48)* (F98) to a cytostatic temperature. Promisingly, all rats with tumors that were treated at their cytostatic hypothermia (20°C for F98 and 25°C for U-87 MG) fully survived the study period. Rats that grew a tumor mass due to insufficient treatment, grew it below the probe (Fig. 4C and Fig. 5C), corroborating our computational analyses suggesting cooling was less effective in the z-axis (Fig. 3C). Fortunately, this problem can be resolved with improved device engineering and is not a biological limitation. Ultimately, cytostatic hypothermia leverages fundamental physics that influences biology broadly. Thus, the effects observed in rats hold better translational promise in larger species (e.g., pigs and humans) compared to conventional species-specific targeted and molecular therapies.

A remaining translational task is development of a patient-centric cytostatic hypothermia device, iterations of which our team has begun designing and patented *(49)*. Permanent neural implants have been clinically available for multiple neurological disorders for years *(50)*.Simultaneously, an FDA-approved device for glioblastoma now exists (worn on the scalp, powered by a backpack battery, and applies tumor-treating fields) *(8)*. This is despite its mild benefit on survival in patients *(8)* and in rats (5.8 ± 2.7 wks median and 10% total survival) *(55)*. To induce hypothermia, heat must be transported out of the tissue. In the middle 20^th^ century, this was attempted through a cryoprobe tethered to an external refrigerator *(14)*. Today, variations of this approach exist, some of which use Peltier plates *(19, 21, 44, 51–53)*. However, current approaches use percutaneous and external device components for heat removal; these increase infection susceptibility (in patients that may be immunocompromised). Similarly, to prove the concept of cytostatic hypothermia in our rodent studies, our tool used an external heat sink and fan. However, a patient-centric device would need to be fully implantable; one such iteration incorporates a more efficient fluid-based method of heat transfer while leveraging skin as a large heat sink (Fig 7A). To enable MRI-compatibility for regular tumor monitoring, a piezoelectric or electromagnetic pump would move the fluid. At the intracranial interface, the rodent studies used a single probe; this was effective but, per the computational analyses (Fig. 4B), it created a temperature gradient. At a larger scale, this gradient could steepen to the point of tissue ablation around the probe. Instead, a multi-probe strategy may deliver cytostatic hypothermia *homogenously* in a larger volume of tissue with minimal tissue displacement (Fig. 7A, inset). However, these designs require computational studies with robust validation. Implantation and design strategies also need to consider the bed and potential CSF cavity after surgical tumor resection. Once a device is implanted, numerous treatment strategies could be explored including lifelong, concomitant, or intermittent hypothermia (as tumors grow more slowly *in vivo (54)*, a therapeutic device may need less than 18 h/d, Fig. 1C). While the tool we engineered can demonstrate the potential of cytostatic hypothermia, developing a patient-centric device may not be too far.

**Figure 7:**
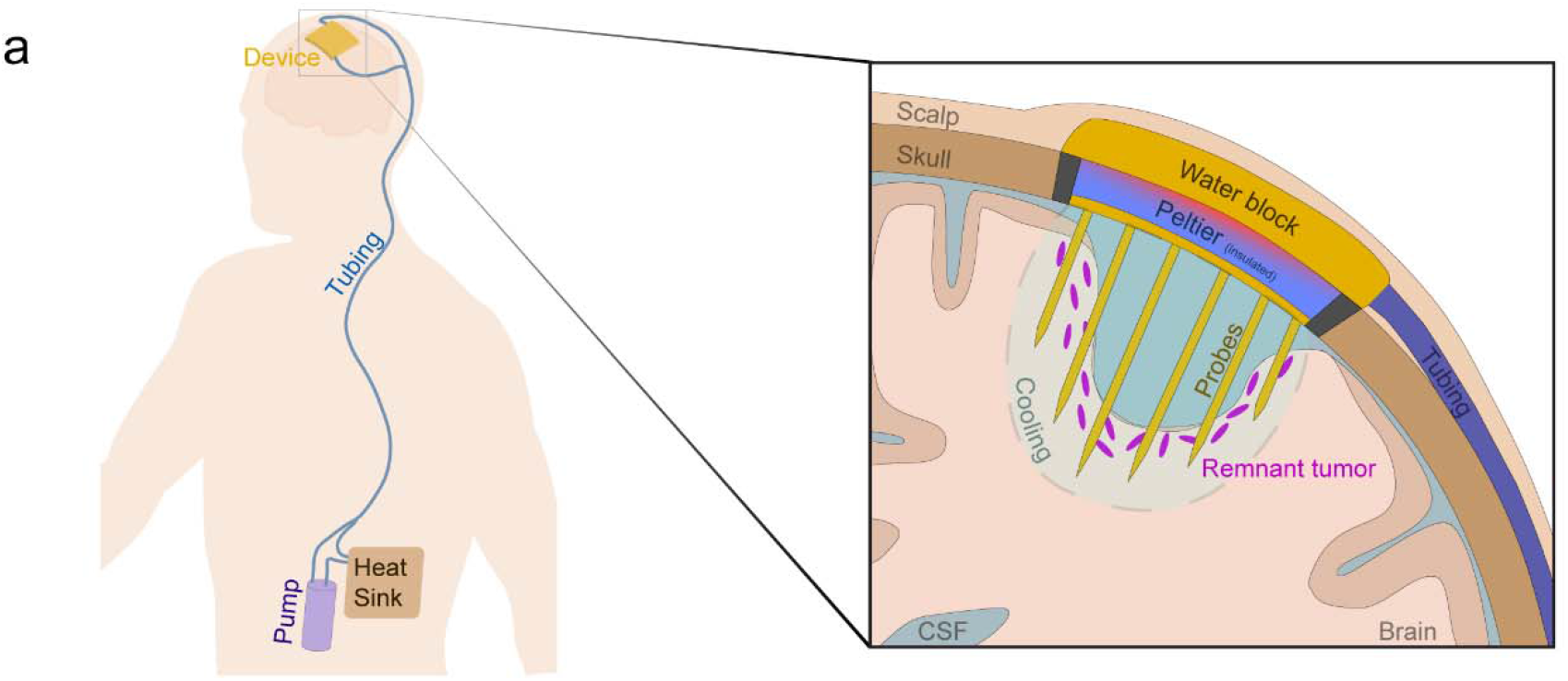
Conceptual model of a translational device to deliver cytostatic hypothermia for GBM. **(A)** Representative diagram of a patient with GBM implanted with an envisioned cytostatic hypothermia system with zoomed inset of device, right. The envisioned system consists of an intracranial device consisting of a multiprobe array in conjunction with a cooling Peltier plate and a water block. An implanted, non-local, non-magnetic pump facilitates water flow through tunneled, subcutaneous tubing to carry heat away from the hot side of the Peltier plate and distribute it to the surrounding skin via the tubing and a subcutaneous heat sink. Inset shows a slice of the cranial cavity after tumor resection with a hypothetical cytostatic hypothermia device and remaining non-resectable tumor cells. A multiprobe array is envisioned to facilitate homogenous cooling at the site of possible tumor recurrence.

Overall, we have provided evidence for the utility of cytostatic hypothermia for slowing GBM growth and demonstrated proof-of-concept *in vivo*. The data, indicating that it affects multiple cellular pathways simultaneously and slows cell division, suggests the opportunities for tumor evolution could be reduced and the likelihood of translating to larger species more likely. In addition, our preliminary computational and *in vivo* studies, in conjunction with our designs and the field of neural interfaces, make a patient-centric device very tangible. As for the biology of cytostatic hypothermia, the field is open for multiple questions and mechanisms that can justify further studies (including those of non-cerebral organs *(23–27)*). Taken together, cytostatic hypothermia could be a novel approach to cancer therapy and eventually serve as an addition to the few options patients with GBM currently have.

## MATERIALS AND METHODS

### a. Cell Culture

All cell lines were purchased from either ATCC or the Cell Culture Facility at Duke. To simplify culturing and passaging, all cells were progressively adapted to a unified medium: Dulbecco’s MEM (Corning 10-013-CV) +NEAA (Gibco) + 10% FBS (Gemini Bio). For metabolite assays, DMEM (Corning 17-207-CV) was supplemented with 5mM Glucose (Gibco), 2 mM Glutamine (Gibco), 1 mM Na^+^ Pyruvate (Gibco), and 10% dialyzed FBS (Gemini Bio 100-108). These media might affect growth rates for individual lines and thus future studies may scrutinize growth under different media formulations under cytostatic hypothermia. Cell lines were passaged twice after thawing for recovery and then used within 5 passages for all experiments. To dislodge cells, 0.05% trypsin (Corning 25-052) was used for U-87 MG and F98, while 0.25% trypsin (Corning 25-053) was used for LN-229 and T98G. Cells were counted using an automated cell counter (Countess II, Invitrogen) calibrated to manual cell counts for each cell line. Cells were plated at a density ranging from 1,000–20,000 cells/well in clear-bottom 96-well plates (Falcon 353219) depending on the desired confluence at the start of the assay. Plated cells were always incubated overnight at 37°C to ensure adherence prior to any experiment. A HeraCell CO2 incubator was used for all experiments at 5% CO2 and set at either 37, 30, 25, or 20°C. Media was carefully replaced every 2–4 days. To prevent losing cells from plates, multichannel pipettes were used to suction media instead of a vacuum.

### b. Imaging and analysis

Imaging was performed on a live-cell microscope (DMi8, Leica Microsystems) with a built-in incubator and CO2 regulator. Well plate dimensions were added to the microscope software (LASX) to enable tile scanning. Images were taken at 5X with a 2×2 field with 25% overlap in 96-well plates. For lengthy imaging periods (>10 min/plate), the incubator and CO2 regulator were used. For analysis, images were merged without stitching as low cell counts prevented accurate automatic stitching. Images were then analyzed through a custom automated ImageJ script to quantify the area coverage of a well plate. To obtain cell morphology (circularity and average size), the script was modified to analyze single cells. For fluorescent live/dead cell detection, an EarlyTox Live/Dead Assay kit was used (Molecular Devices) and the microplate was imaged with a laser and GFP and TXR filters. For immunotherapy experiments, images were taken at 10X with a 4×4 field and a laser through a GFP filter. The ImageJ script was modified to count the number of GFP^+^ cells. All analyzed images were then processed through custom Python scripts to organize the data for analysis.

### c. Molecular assays

Cell cycle analyses were conducted with a Propidium Iodide Flow Cytometry kit (ab139418, Abcam). Samples were run on a flow cytometer (Novocyte 2060, ACEA Biosciences). Data were analyzed on FlowJo v10.7.

Intracellular ATP was detected from homogenized cells via a Luminescent ATP Detection Assay kit (ab113849, Abcam). Luminescence was detected in a microplate reader (SpectraMax i3x, Molecular Devices).

Metabolites were detected from media through luminescent based assays. Cells were grown in four plates with modified media at 37°C overnight and then media was replaced. All plates were then left for 3 days at 37°C after which media was collected and stored at −80°C from one plate and replaced in the others. Three plates were then moved to either 30, 25, or 20°C. Media was collected, stored at −80°C, and replaced every 3 days. Images were taken in a livecell microscope prior to collecting media. Glucose consumption (Glucose-Glo, J6021, Promega), Lactate production (Lactate-Glo, J5021, Promega), Glutamine consumption (Glutamine/Glutamate-Glo, J8021, Promega), and Glutamate production (Glutamate-Glo, J7021, Promega) were assayed from diluted samples. Luminescence was detected in the microplate reader and calibrated to a standard curve of each metabolite. Data were normalized to well coverage area quantified from images and a custom ImageJ macro.

For cytokine analysis, cells seeded in 24-well plates at different densities for the 37°C and 25°C plates to reach similar confluence at the time of the assay. All plates were incubated overnight at 37°C and then incubated for 3 days at their respective temperature. Media was then collected and stored at −80°C from one plate. The media in the remaining plates were replaced and the plates were moved to 25°C. Media was then collected, stored, and replaced every 3–4 days. Cytokines were detected from media samples with a custom multi-plex ELISA kit (LEGENDplex, BioLegend) and our flow cytometer (Novocyte 2060).

### d. Adjuvant studies

Chemotherapy: Temozolomide (Sigma Aldrich) was dissolved in DMSO at 20 mg/mL and stored at −20°C in aliquots. Aliquots were used within 2 months. For these assays, cell lines were grown in the microplate overnight and then either kept at 37°C or moved to 25°C. TMZ at one of 3 doses (0, 500, or 1000 μM) was added to the 37°C plate and incubated for 4 days. The 0 μM TMZ dose consists of DMSO at an equivalent concentration to that in the 1000 μM TMZ group. The 25°C plate was first incubated for 2 days at 25°C (cytostatic for the three lines), then TMZ was added at one of the three doses, and the plate was then further incubated at 25°C for 4 days. After 4 days of TMZ treatment, the media was completely replaced in both plates, and the plates were moved back to 37°C with daily imaging in a live-cell microscope. The line graphs of Figure 3a–c are overlayed to be able to easily compare TMZ treatment at 37°C vs. TMZ treatment at 25°C.

Immunotherapy: Briefly, using a previously described protocol *(55)*, CAR T cells were generated by harvesting splenocytes from mice, and transducing them with a retrovirus that induces CAR T expression specifically targeting EGFRvIII on CT2A tumor cells. CAR T cells were stored in liquid nitrogen 4 days post-transduction. All experiments used CAR T cells 5–7 days post-transduction. For immune killing assays, 2000 EGFRvIII^+^ GFP^+^ CT2A cells were grown per well overnight in modified DMEM media as described previously. In pre-treatment studies, these plates were incubated for 4 or 7 days at 25°C. For experiments, different dosages of CAR T cells were added the following day with phenol red-free RPMI media and microplates were incubated at either 37 or 25°C. Microplates were imaged at regular intervals as described previously. The number of GFP^+^ cells was quantified through an ImageJ macro.

### e. Device fabrication

Device design and manufacturing was done in collaboration with the Pratt Bio-Medical Machine Shop at Duke University. Designs were developed on MasterCam and Fusion360 (fig. S6). Most materials and parts were obtained through McMaster, or vendors such as Digi-Key, Newark Element, Mouser, or Amazon. All components and manufacturing/fabrication methods are described in the Supplementary Materials and Methods.

### f. Caging and treatment set-up

To enable the free movement of awake rats under treatment, a custom caging and treatment setup was developed.

#### Cage and lever-arm

Rat housing was constructed by flipping 22qt plastic containers (Cambro) which were made hollow by sawing off the closed end. They were then sealed onto ¼”x15”x15” acrylic sheets (TAP Plastics) with cement for plastic (7515A11, McMaster). Parts were 3D printed to hold a water bottle and a lever arm. The two-axis lever arm consisted of 3D printed parts holding ¼” steel rods (McMaster) and a custom-made counterweight.

#### Electrical/treatment

Patch cables were made with male connectors (GHR-06V-S, JST Sales America) on either end of 12” jumper cables (AGHGH28K305, JST Sales America). These were protected by a spring (9665K84, McMaster) and the joints strengthened with epoxy and hot glue. A slipring (736, Adafruit Industries), to enable free rotation, had a female connector soldered and epoxied to an end hovering inside the cage. The patch cable connected the slipring to the Cooler.

Outside the cage, a variable voltage power supply (HM305, Hanmatek) was connected to the slipring with alligator clips. Each one powered the Peltier plate in one Cooler. USB 5V adapter towers (Amazon) were used to power the fans of multiple Coolers and connected with a USB jack on one end and alligator clips to the slipring on the other. Arduinos (Uno Rev 3, Arduino) connected to a laptop (through USB adapter towers) were connected to sliprings through a voltage-divider circuit on a breadboard and alligator clips. One Arduino was used per device for temperature monitoring.

### g. Animals

All animal procedures were approved by the Duke IACUC. Fischer (CDF) and Nude (RNU) rats were purchased from Charles River at 7–9 weeks of age. All procedures began at 8–10 weeks of age. Tumor inoculation, device implantation, Cooler attachment, monitoring and maintenance, and euthanasia are described in detail in the Supplementary Materials and Methods.

### h. MRI

All MRI was performed in the Center for In Vivo Microscopy (CIVM) at Duke University. For images after implantation, the Cooler was unscrewed and removed from the Interface under anesthesia. Rats were positioned on a custom 3D-printed bed and eye gel and ear covers were applied. The rats were imaged in a Bruker BioSpin 7T MRI machine (Bruker, Billerica MA) with a 4-element rat brain surface coil. T1 weighted FLASH (TE/TR = 4.5/900 ms; Flip angle = 30°; 3 averages) and T2 weighted TurboRARE (TE/TR = 45/8000 ms; RARE factor of 8; 2–3 averages) axial images were acquired at a resolution of 100 μm in-plane and slice thickness of 500 μm. During all protocols, breathing was monitored via telemetry. Animal body temperature was maintained with warm air circulation in the bore of the magnet. Upon completion, if an Interface was present, the Cooler was reattached with thermal paste applied between the copper contacts. T2-weighted images were analyzed on 3D Slicer v4.10.2 (www.slicer.org) with assistance of animal imaging experts at the CIVM for tumor-take confirmation and volume quantification. Volume analysis was conducted by manually tracing tumor boundaries on each MR slice and volume was computed by the software.

### i. Histology

Brains were cryo-sectioned into 12 μm slices, collected on Superfrost+ slides, and the slides were stored at −20°C. Some slides were stained with Hematoxylin & Eosin (H&E) and imaged under our microscope with a color filter. Other slides were stained with immunohistochemistry markers (from Abcam) and imaged under our microscope with fluorescent lasers. These included: mouse anti-GFP (ab1218), mouse anti-human Ku80 (ab119935), rabbit anti-rat NeuN (ab177487), chicken anti-rat GFAP (ab4674), rabbit anti-rat CD45 (ab10558), rabbit anti-Ki-67 (ab16667), and rabbit anti-rat/human cleaved caspase 3 (ab49822). Secondary antibodies included: Goat anti-mouse Alexa Fluor 488 (ab150113), Goat anti-rabbit Alexa Fluor 594 (ab150080), and Goat anti-chicken Cy5.5 (ab97148). Briefly, slides were washed with PBS and 0.5% Triton, blocked with 4% goat serum for 2 hours, followed by staining with primary antibodies diluted in 1% BSA, overnight at 4°C. Secondary-only controls did not receive any primary antibody. The next day, slides were washed with PBS, stained with secondary antibodies diluted in 1% BSA for 2 hours, counterstained with DAPI for 15 minutes, washed and coverslipped with Fluoromount G (Southern Biotech), and left to dry overnight at room temperature. The subsequent day, nail polish was applied to the edges of the slides and then they were stored in boxes at room temperature.

Histological images were acquired by either a color camera (for H&E) or a fluorescence camera (for IHC) attached to the DMi8 microscope previously described. Whole-section tile-scanned images were taken at 10X zoom with 25% overlap and automatically stitched. Care was taken to ensure imaging for any stain group was conducted in a single sitting. Images were then processed via a custom ImageJ script to create composites with suppressed background signal. The final images and H&E slides were then independently analyzed by an expert neuropathologist who was blinded to the experimental conditions.

### j. Computational modelling

Modelling was performed on COMSOL v5.2 to study heat transfer between the probe and surrounding tissue via the finite element method. Both the steady state temperature and transient effects of initiating and stopping cooling were studied. The geometry of the model consisted of a rat brain model containing a spherical volume of tissue (which could be interpreted as both a bulk tumor or the extent of a small, infiltrating tumor) and gold probe. The rat brain model, acquired from the literature *(30)*, was generated from MRI data and smoothened to reduce anatomical indentations. The probe was modeled as a cylinder with a diameter of 1mm and height of 2.5 mm with a 45° chamfer at the tip. The tip of the probe was embedded in the tumor which was modeled as a sphere with diameters of 2 mm and 3 mm. Heat transfer was modeled with the Pennes’ bioheat equation. A negative heat flux was applied to the top face of the probe to cool the probe. All other outer surfaces were fully insulated with zero heat flux.

All tissues and materials used in the model were assumed to be homogeneous and isotropic – features such as brain blood vessels were not modeled. Multiple parameters used for the tissues, perfusion, and materials were investigated (table S5). Original perfusion values in units of ml/100□g/min were converted into s^-1^ using a blood density of 1050 kg/m^3^.

The blood perfusion and metabolic heat generation parameters were temperature-dependent based on the following equations:

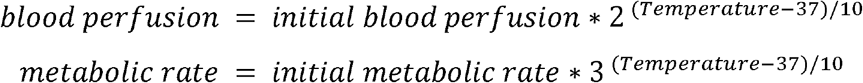

### k. Statistical analysis

All statistical analyses were performed on GraphPad Prism v9.0.2. Error is expressed as standard deviation (SD). Where appropriate, one-way, repeated measures, and two-way ANOVAs were performed with Dunnett’s multiple comparisons post-hoc test unless otherwise specified. Survival was analyzed with the Log-rank Mantel-Cox test. Significance was set to 0.05 for all studies.

## List of Supplementary Materials

Materials and Methods

Fig S1 to S11

Tables S1 to S7 (Excel files)

Movies S1 to S2

## Acknowledgments

We are grateful to: Dr. Nalini Mehta and Sheridan Carroll for generating CAR T cells; the Center for In Vivo Microscopy at Duke University for MR imaging; the staff (Kaylee Lynn, Bridget Pickle, Jesse DeGraff, and Fernando Orozco) and veterinarians (Drs. Felicitas Smith and Clay Rouse) at the Duke Vivarium for their assistance in setting up and maintaining animal experiments; Edward Mallon for his advice on setting up a temperature recording system; and Dr. Syed Ather Enam for his neurosurgical insights.

## Author contributions

S.F.E. conceived the project. S.F.E., C.Y.K., J.H., C.S.T., and E.I. conducted the in vitro studies and data analysis. S.F.E., B.J.K, and M.I.B. conducted the in vivo studies and histology. R.C. and S.F.E. conducted the computational studies. S.J.B. conducted MR imaging and analysis. A.F.B. and S.F.E. conducted histological analysis. S.F.E. and S.J.O. designed and fabricated the devices. S.F.E. wrote the manuscript. All authors edited and reviewed the manuscript. S.F.E., J.G.L., and R.V.B. supervised the studies. R.V.B. funded all the studies.

## Data and materials availability

All data and code that support and produce the findings of this study are available from the corresponding authors upon request.

---

**Supplementary Figure 1:**
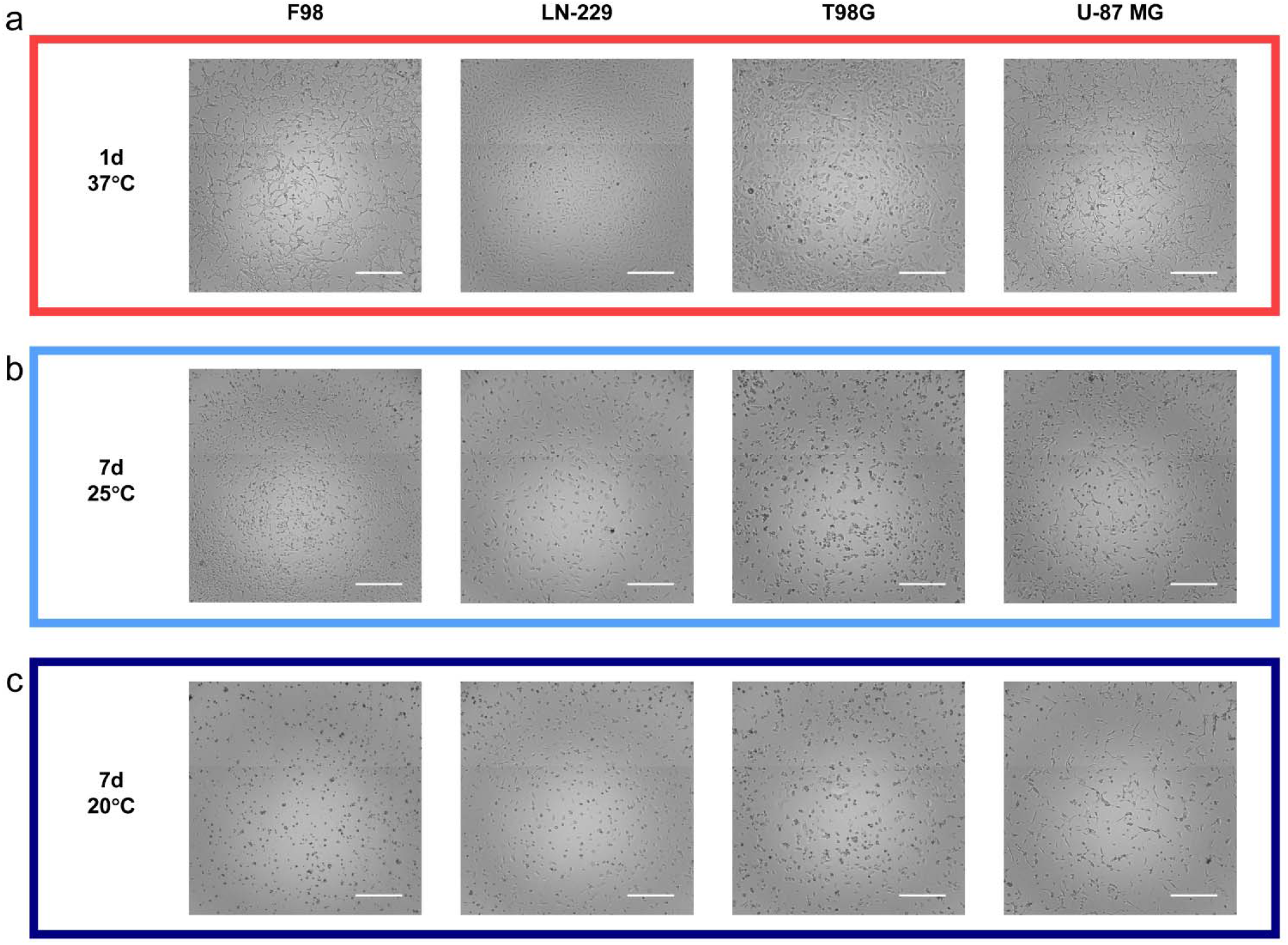
Brightfield images of GBM lines under different temperature conditions. All cells received overnight incubation at 37°C and then were either left at 37°C or transferred to 25°C or 20°C. White scale bars = 500 μm. **(A)** Images of cells after 1 day at 37°C. **(B)** Images of cells after 7 days at 25°C. **(C)** Images of cells after 7 days at 20°C.

**Supplementary Figure 2:**
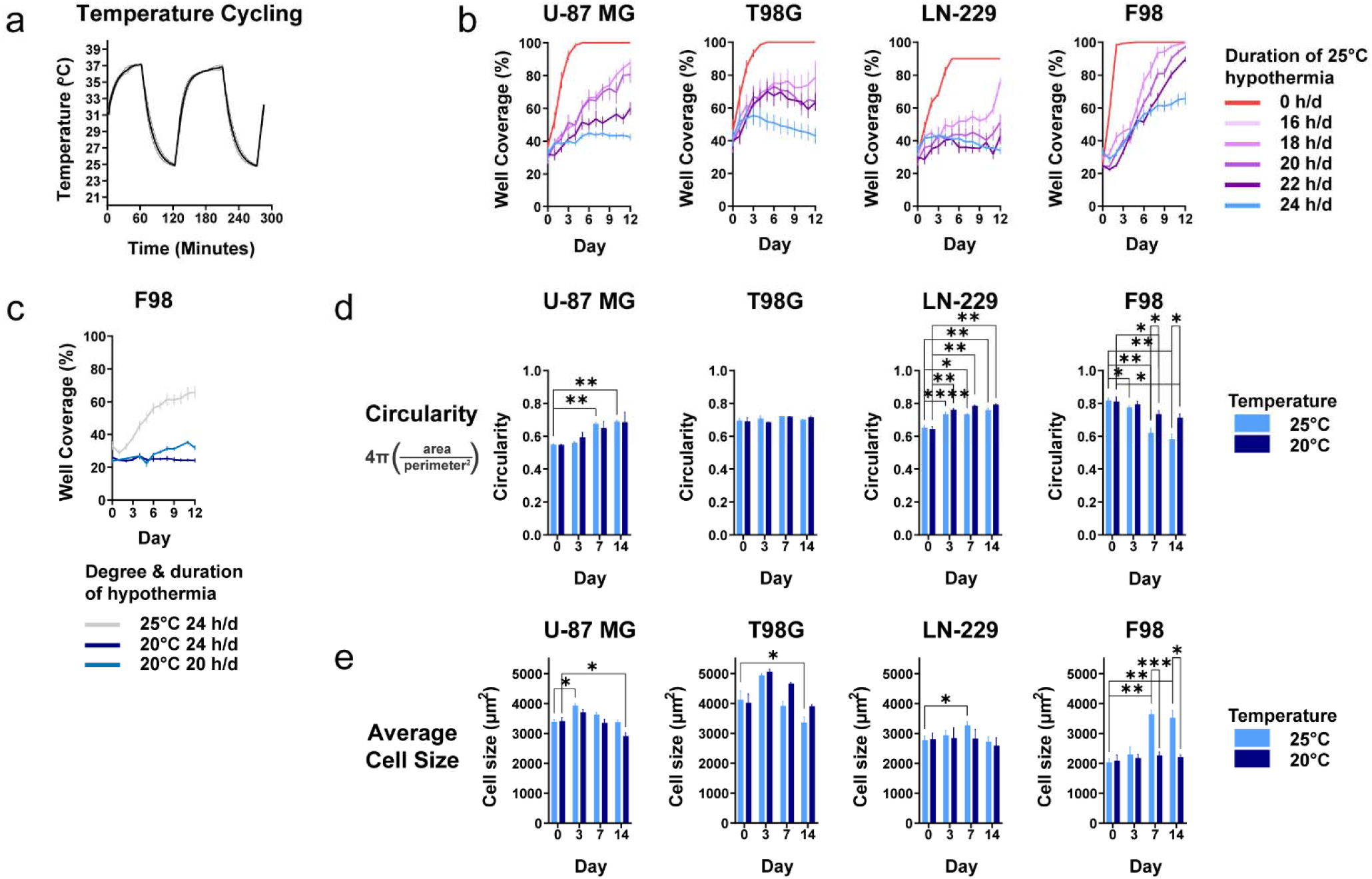
Effect of intermittent hypothermia on cell growth and continuous hypothermia on cell morphology. **(A)** Plot demonstrating the change of media temperature when a microplate is transferred between 37°C and 25°C cell incubators (n = 2). Thermistors were attached on the bottom of a central well and lateral well. Resistance was measured with a voltage-divider circuit and Arduino and converted to temperature with a script. **(B)** GBM growth rates under intermittent 25°C hypothermia without 25°C pre-treatment (n = 8). Plates were transferred between 37°C and 25°C incubator for varying hours/day (h/d). **(C)** F98 growth rate under continuous 25°C or 20°C hypothermia or 20 h/d intermittent 20°C hypothermia with 4 days of 20°C pretreatment. **(D) and (E)** Average tumor cell circularity and average size under hypothermia (n = 8). Circularity and average size were determined through ImageJ. Two-way ANOVAs were conducted with Dunnet’s multiple comparison tests post-hoc (\**p*<0.05, \*\**p*<0.01, \*\*\**p*<0.001, \*\*\*\**p*<0.0001). Specific adjusted *p*-values are provided in Supplementary Tables 2 and 3. All graphs show mean ± standard deviation.

**Supplementary Figure 3:**
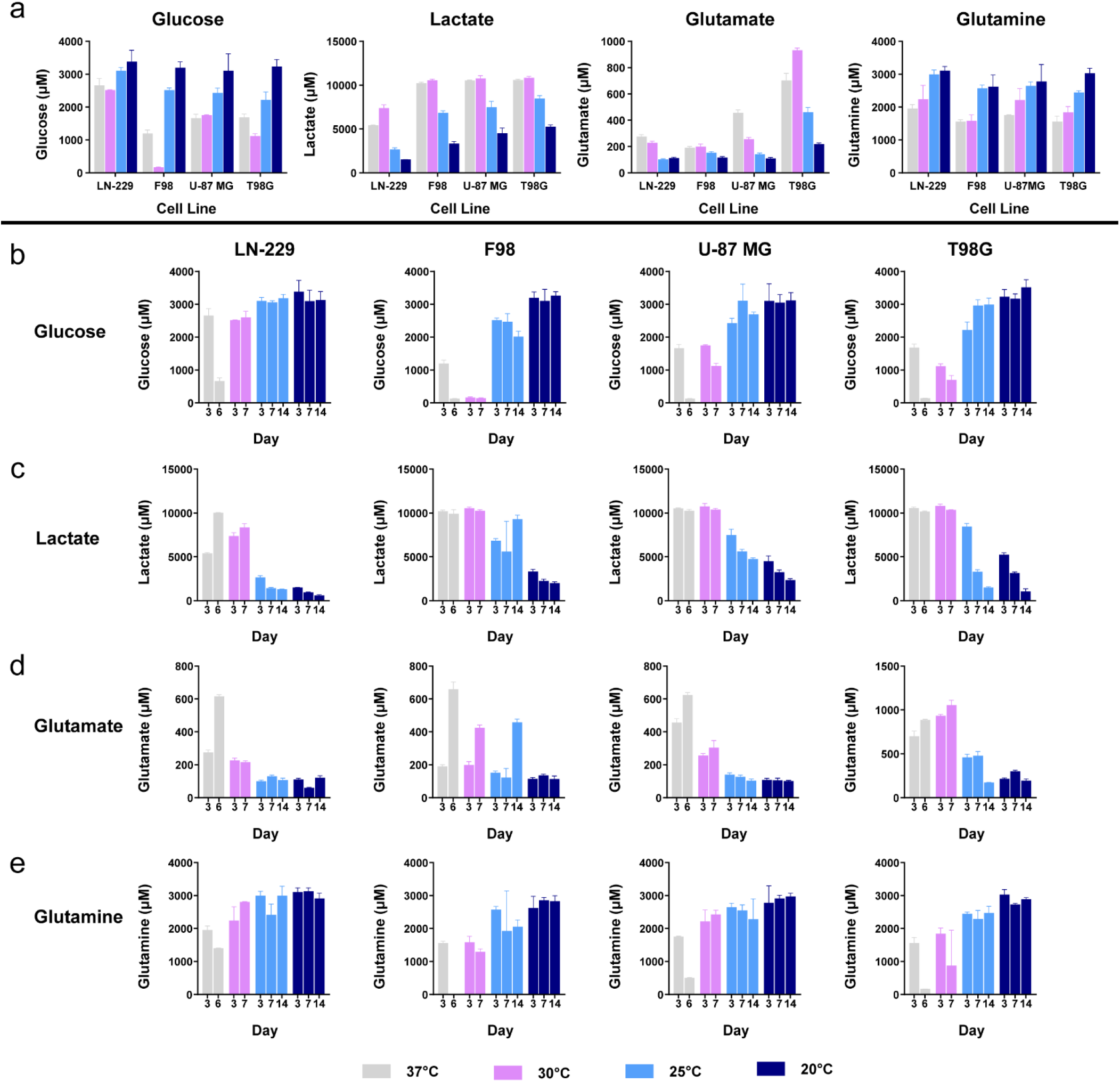
Metabolite production and consumption under hypothermia. **(A)** Metabolite levels in media after 3 days at 37°C or 3 days at hypothermic temperatures (30, 25, or 20°C) (n = 3). **(B)** Glucose levels in media at 3, 7, and 14 days at different temperatures. **(C)** Lactate levels in media at 3, 7, and 14 days at different temperatures. **(D)** Glutamate levels in media at 3, 7, and 14 days at different temperatures. **(E)** Glutamine levels in media at 3, 7, and 14 days at different temperatures. All graphs show mean ± standard deviation.

**Supplementary Figure 4:**
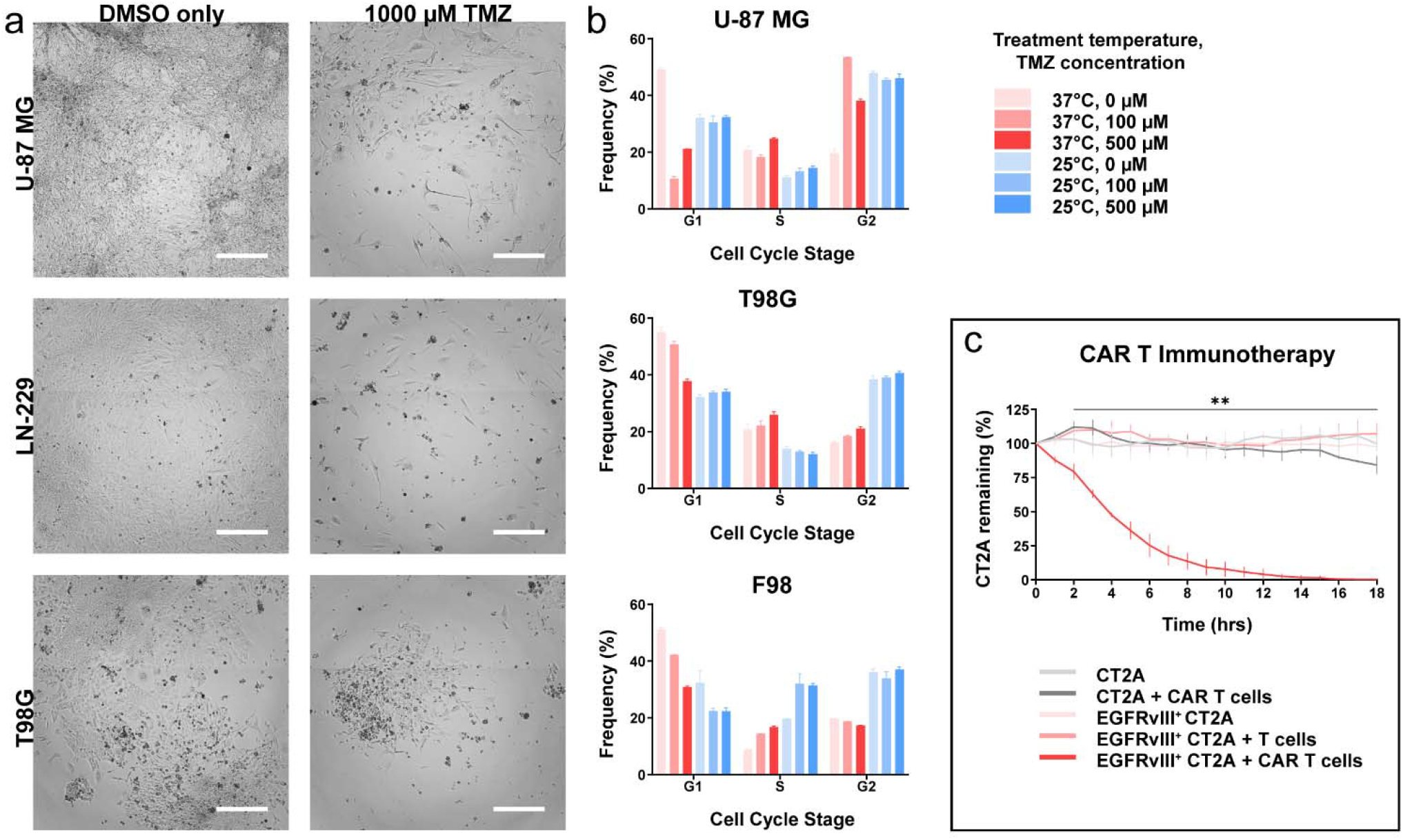
Effect of cytostatic hypothermia on TMZ and validation of CAR T cells. **(A)** Brightfield images representing U-87 MG, LN-229, and T98G pre-treated with DMSO only or 1000 μM TMZ in DMSO, both under hypothermia. These images were taken 12 days after TMZ was removed and the cells had been returned to incubate at 37°C. White scale bar = 500 μm **(B)** Effect of TMZ and hypothermia on cell cycle progression (n = 3). U-87 MG (above), T98G (middle), or F98 (bottom) were assayed immediately after TMZ treatment under 37°C or 25°C. **(C)** Graph demonstrating the specific killing of EGFRvIII^*+*^ CT2A cells by CAR T cells only (n = 4). Repeated measures ANOVA was conducted with Dunnett’s multiple comparisons test comparing the group of EGFRvIII^*+*^ CT2A cells with CAR T cells to EGFRvIII^-^ CT2A only (\*\**p* < 0.01 at least in all comparisons).

**Supplementary Figure 5:**
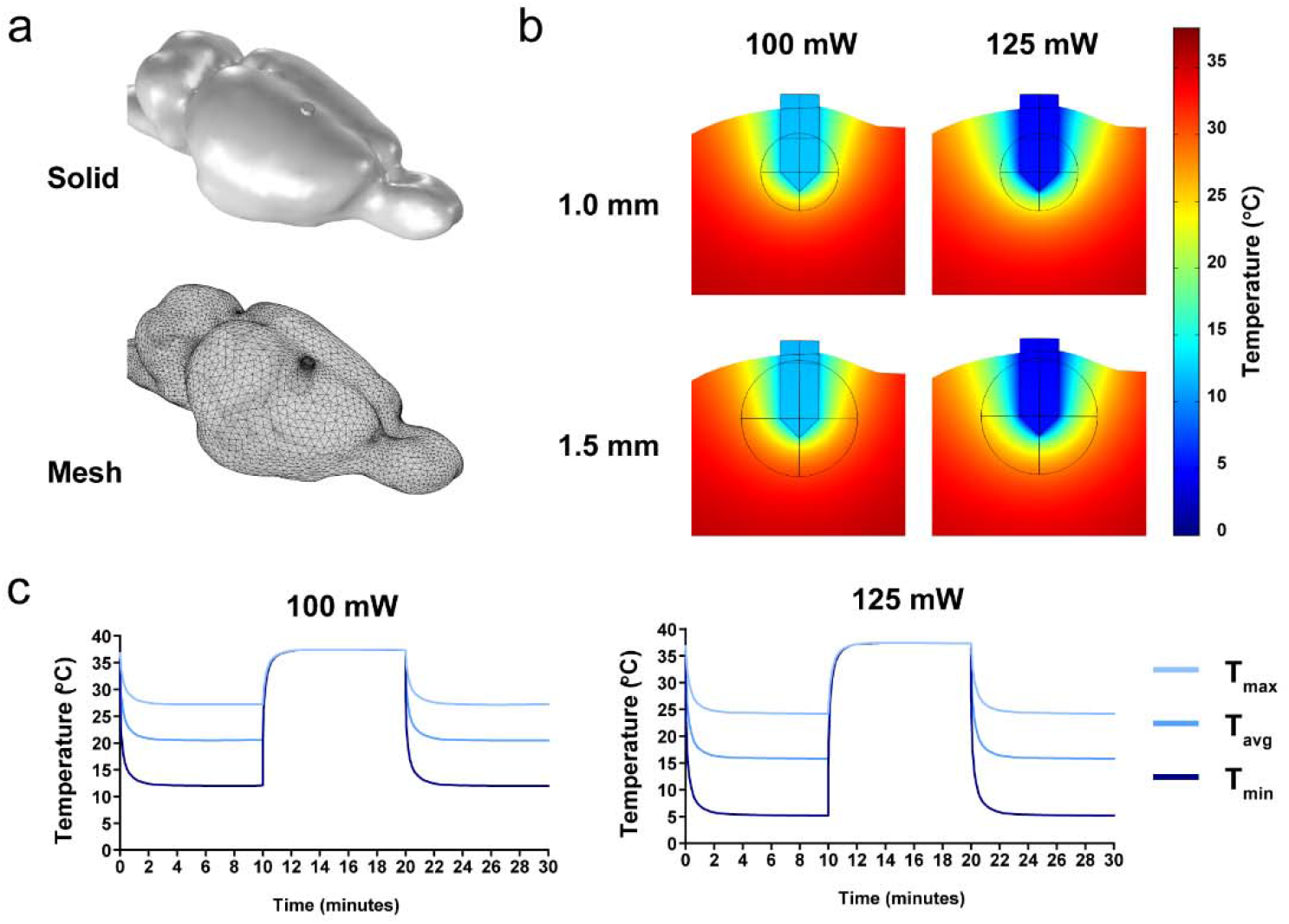
Finite-element analysis of local intratumoral hypothermia. **(A)** Above: Smoothened rat brain model obtained from the literature with the addition of a probe and tumor. Below: fine mesh added to the model for finite-element analysis. **(B)** Coronal slices depicting degree and extent of hypothermia depending on tumor size (1 mm and 1.5 mm radius) and heat energy pulled (100 and 125 mW). **(C)** Time-dependent study modeling maximum, average, and minimum tumor temperatures over time. Cooling was begun at t = 0, stopped at t = 10, and restarted at t = 20 minutes.

**Supplementary Figure 6:**
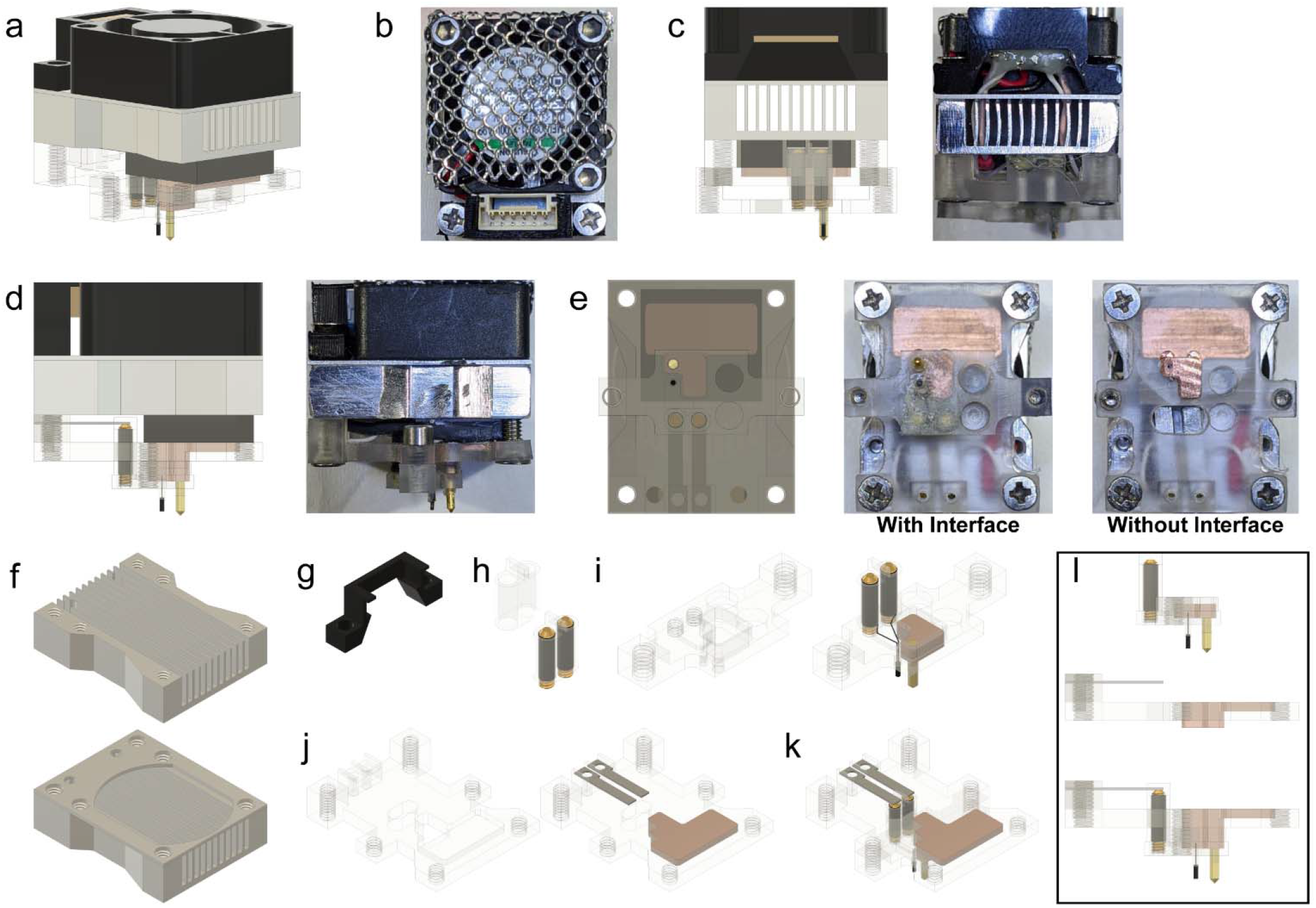
Local hypothermia rodent device. **(A)** Rendered front angle view of device. **(B)** Top view of device. Aluminum mesh is visible above a fan. Beige connector is visible surrounded by a 3D printed cover. **(C)** Rear view of device. Left: render. Right: picture shows wires terminating at connector where they were soldered and protected with epoxy. **(D)** Side view of device. All components are visible with this perspective. **(E)** Bottom view of device. Left: render. Middle: Picture including the presence of Interface. Right: Picture without the Interface. Steel shims, Peltier wires, and copper part are visible through the polycarbonate base. **(F)** Render of heat sink without cover (above) and with cover (below). Holes were drilled to enable passage of wires and fastening of screws. **(G)** Render of 3D-printed component to secure the beige connector. **(H)** Render of polycarbonate part to protect the heat-shrink covered brass screws. **(I)** Render of polycarbonate Interface base alone (left) and with all parts (right). Thermistor wires are wrapped around the brass screws under the heat shrink. Copper part with gold needle is pressed into the polycarbonate base. **(J)** Render of polycarbonate plate of Cooler alone (left) and with all parts (right). Steel shims are components of the thermistor circuit. **(K) and (L)** Renders of Interface and polycarbonate plate of Cooler as they are contacted together to enable heat conduction and closing the thermistor circuit.

**Supplementary Figure 7:**
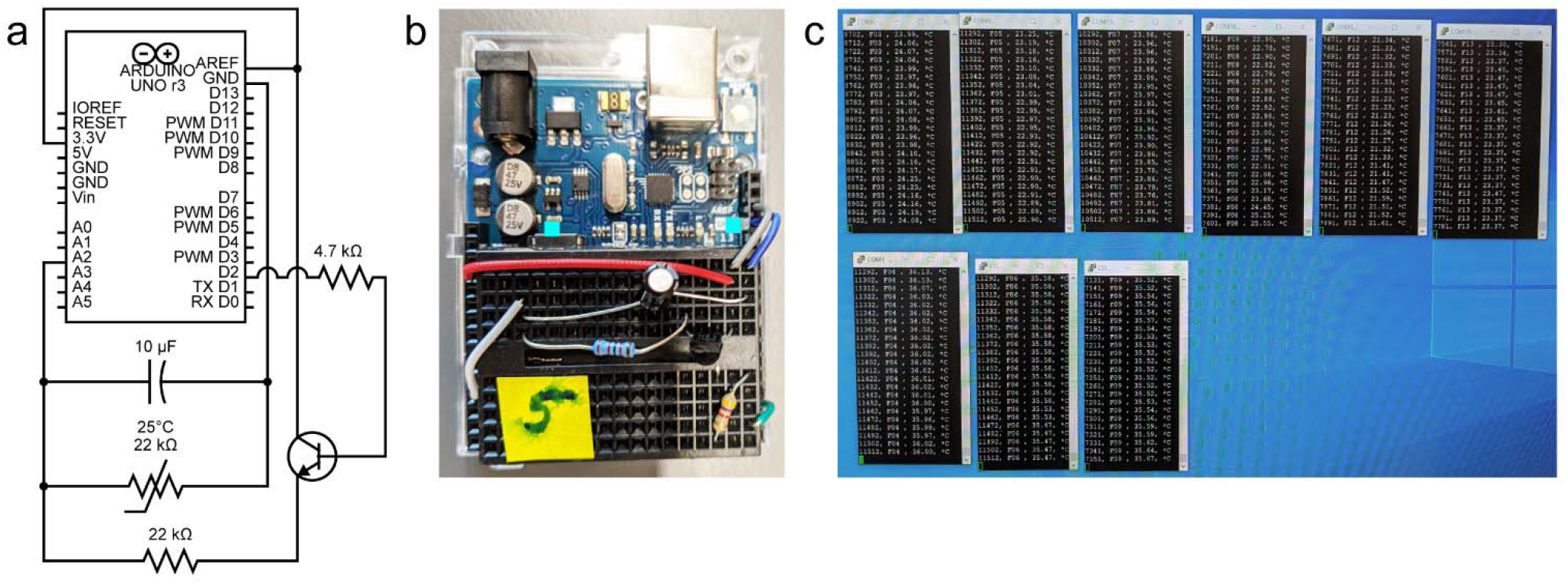
Thermistor circuit for temperature monitoring. **(A)** Schematic demonstrating the voltage-divider circuit used to measure resistance across the thermistor. A digital pin from the Arduino was used to take intermittent and averaged recordings (every 10s). **(B)** Photo of voltage-divider circuit on a breadboard above the Arduino. One of these was used per rat for temperature monitoring. Arduino code converted the resistance across the thermistor to temperature using a manufacturer supplied beta-value. **(C)** Picture of computer screen depicting continuous temperature monitoring through PuTTY. The computer was also connected to the network to enable temperature monitoring outside the vivarium.

**Supplementary Figure 8:**
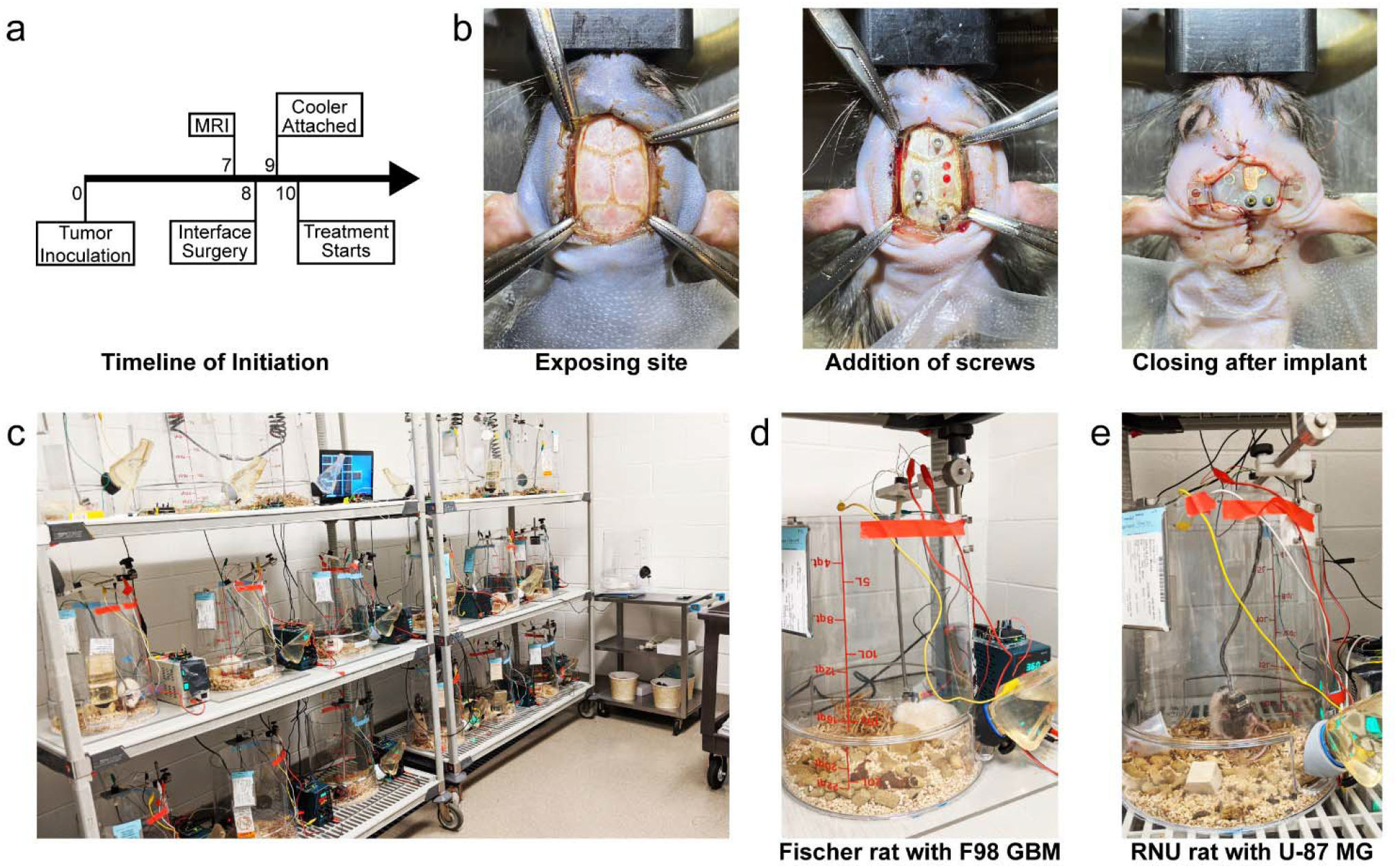
*In vivo* Interface implantation and treatment application. **(A)** Timeline of tumor inoculation, MRI, Interface implant surgery, Cooler attachment, and treatment initiation. **(B)** Photos of implantation surgery in an RNU rat 8 days after tumor inoculation. Left: the surgical site is re-exposed and hemostasis established. Middle: burr holes were drilled for the intratumoral probe, thermistor, and four titanium screws. The titanium screws were tightened by hand. Right: suturing around the Interface which was attached to the skull using dental cement. The copper part for heat conduction and the brass screws are visible for contact with the Cooler. **(C)** Set up of multiple cages, power supplies, temperature monitoring, and a central computer in the vivarium. Rats were weighed in an empty cage visible on the right. **(D)** Photo of a Fischer rat under treatment. Patch cable, slip ring, lever arm, power supply, and water bottle are visible. **(E)** Nude (RNU) rat under treatment.

**Supplementary Figure 9:**
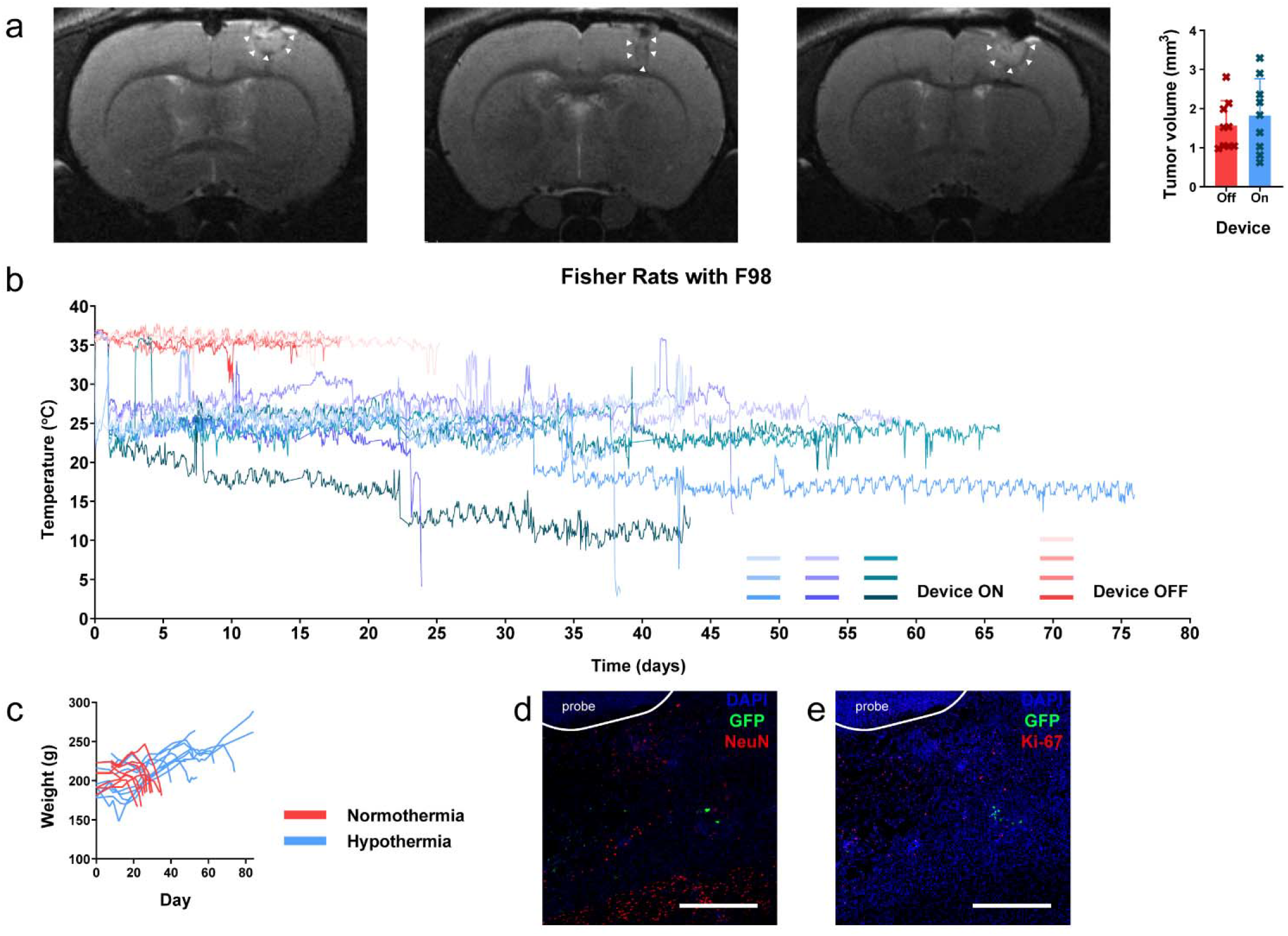
Hypothermia treatment of Fischer rats inoculated with F98 GBM. **(A)** T2-weighted MR images 7 days after tumor inoculation confirming tumor development. White arrows demarcate tumor boundaries. Tumor volumes between the two treatment groups were not significantly different (Unpaired t test) prior to initiating treatment. Graph shows mean ± standard deviation. **(B)** Continuous temperature monitoring of 9 rats with their device switched on and 4 rats with their device switched off. The remaining 5 normothermia rats did not have temperature monitoring. **(C)** Rat weights through the study period. **(D-E)** Histological sections of brain from a surviving rat after hypothermia treatment. White scale bar = 500 μm. **(D)** Presence of GFP^+^ cells (green) below the cooling probe among cells staining for NeuN (red). **(E)** GFP^+^ cells (green) did not colocalize with Ki-67 (red) in the surviving rat.

**Supplementary Figure 10:**
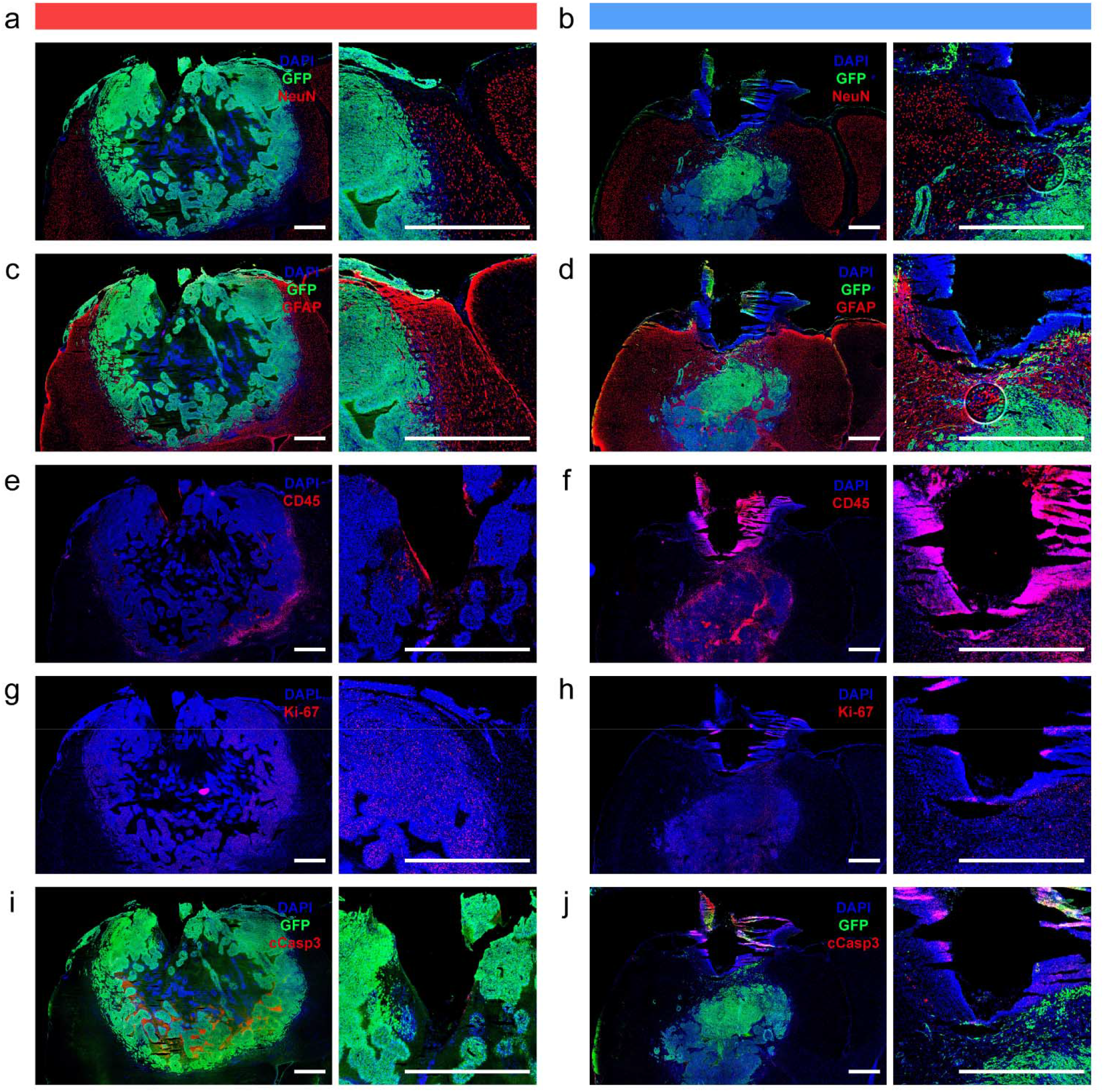
Representative immunohistochemistry of Fischer rats with F98 GBM with and without hypothermia. Red bar indicates rat from the normothermia group that was euthanized at ~4 wks after tumor inoculation; blue bar indicates rat from the hypothermia group that was euthanized at ~8 wks after tumor inoculation. **(A-B)** NeuN staining demonstrating intact neuronal nuclei around the tumor (A) and near the hypothermia probe (B). **(C-D)** GFAP staining demonstrating peri-tumoral and ipsilateral reactive gliosis (C) and intact glial populations near the hypothermia probe (D). **(E-F)** CD45 staining to demonstrate leukocyte inflammation within the tumor (E) and within the immediate region adjacent to the hypothermia probe (F). **(G-H)** Ki-67 staining demonstrating proliferative activity in tumor bulk (G) while activity is evidently reduced adjacent to the probe (H). **(I-J)** Cleaved caspase 3 staining demonstrating extensive tumor-intrinsic apoptosis in the rat from the normothermia group (I), and apoptosis in the immediate region around the cooling probe (J). White scale bars = 1000 μm.

**Supplementary Figure 11:**
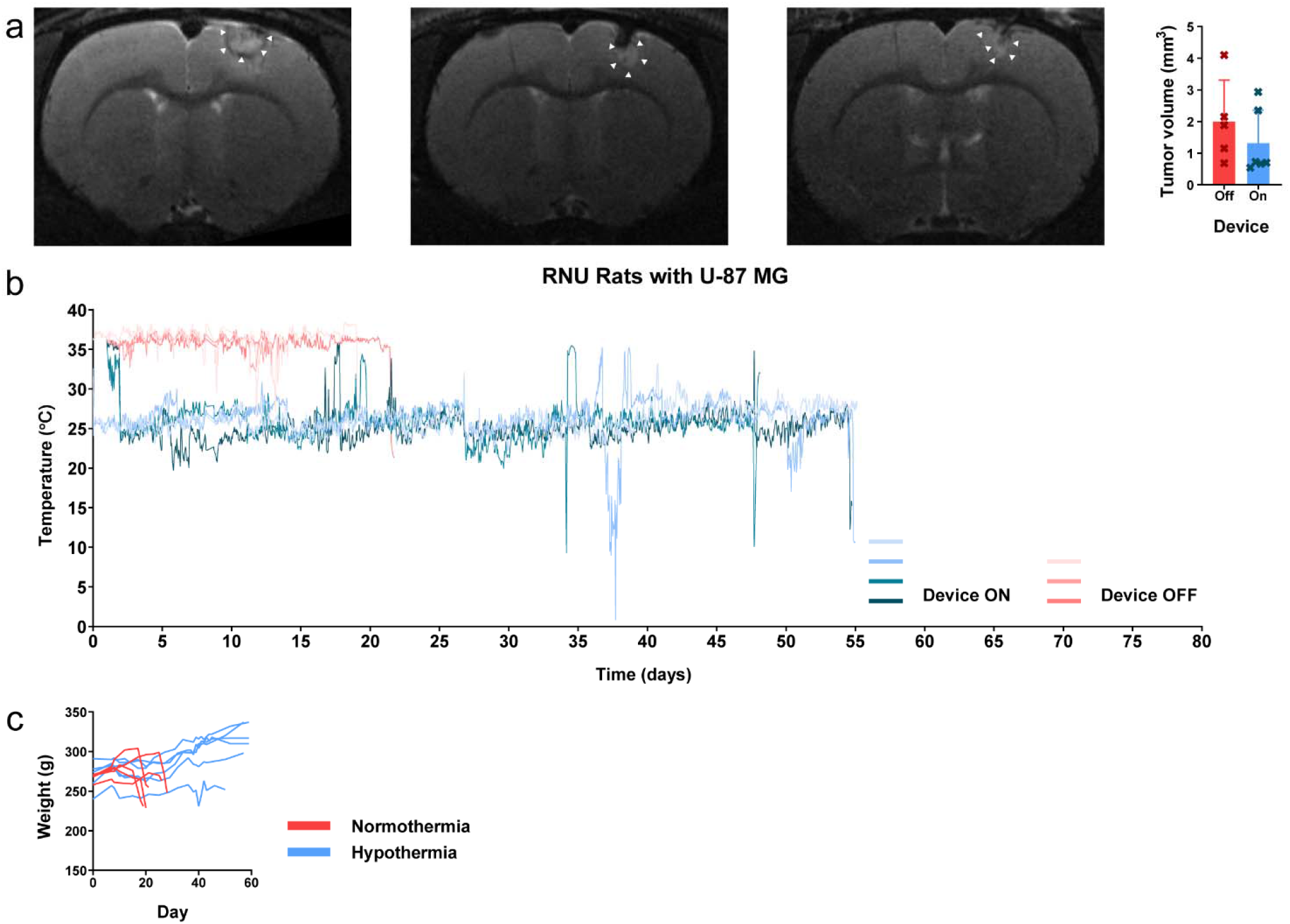
Hypothermia treatment of Nude (RNU) rats inoculated with U-87 MG. **(A)** T2-weighted MR images 7 days after tumor inoculation confirming tumor development. White arrows demarcate tumor boundaries. Tumor volumes between the two treatment groups were not significantly different prior to initiating treatment. Graph shows mean ± standard deviation. **(B)** Continuous temperature monitoring of 4 rats with their device switched on and 3 rats with their device switched off. The remaining 2 treatment rats had premature failure of their thermistors with erratic readings and are thus not displayed. The remaining 2 normothermia rats did not have temperature monitoring. **(C)** Rat weights through the study period.

## References and Notes

1. M. Koshy, J. L. Villano, T. A. Dolecek, A. Howard, U. Mahmood, S. J. Chmura, R. R. Weichselbaum, B. J. McCarthy, Improved survival time trends for glioblastoma using the SEER 17 population-based registries. J. Neurooncol. 107, 207–12 (2012).

2. Q. T. Ostrom, G. Cioffi, H. Gittleman, N. Patil, K. Waite, C. Kruchko, J. S. Barnholtz-Sloan, CBTRUS Statistical Report: Primary Brain and Other Central Nervous System Tumors Diagnosed in the United States in 2012–2016. Neuro. Oncol. 21, v1–v100 (2019).

3. M. Rapp, J. Baernreuther, B. Turowski, H. J. Steiger, M. Sabel, M. A. Kamp, Recurrence Pattern Analysis of Primary Glioblastoma. World Neurosurg. 103, 733–740 (2017).

4. J. P. Kirkpatrick, N. N. Laack, H. A. Shih, V. Gondi, Management of GBM: a problem of local recurrence. J. Neurooncol. 134, 487–493 (2017).

5. A. A. Brandes, A. Tosoni, E. Franceschi, G. Sotti, G. Frezza, P. Amistà, L. Morandi, F. Spagnolli, M. Ermani, Recurrence pattern after temozolomide concomitant with and adjuvant to radiotherapy in newly diagnosed patients with glioblastoma: Correlation with MGMT promoter methylation status. J. Clin. Oncol. 27, 1275–1279 (2009).

6. M. Weller, M. van den Bent, J. C. Tonn, R. Stupp, M. Preusser, E. Cohen-Jonathan-Moyal, R. Henriksson, E. Le Rhun, C. Balana, O. Chinot, M. Bendszus, J. C. Reijneveld, F. Dhermain, P. French, C. Marosi, C. Watts, I. Oberg, G. Pilkington, B. G. Baumert, M. J. B. Taphoorn, M. Hegi, M. Westphal, G. Reifenberger, R. Soffietti, W. Wick, European Association for Neuro-Oncology (EANO) guideline on the diagnosis and treatment of adult astrocytic and oligodendroglial gliomas Lancet Oncol. (2017), doi:10.1016/S1470-2045(17)30194-8.

7. D. Gramatzki, P. Roth, E. J. Rushing, J. Weller, N. Andratschke, S. Hofer, D. Korol, L. Regli, A. Pangalu, M. Pless, J. Oberle, R. Bernays, H. Moch, S. Rohrmann, M. Weller, Bevacizumab may improve quality of life, but not overall survival in glioblastoma: An epidemiological study. Ann. Oncol. 29, 1431–1436 (2018).

8. S. Kesari, Z. Ram, Tumor-treating fields plus chemotherapy versus chemotherapy alone for glioblastoma at first recurrence: a post hoc analysis of the EF-14 trial. CNS Oncol. (2017), doi: 10.2217/cns-2016-0049.

9. A. Jain, M. Betancur, G. D. Patel, C. M. Valmikinathan, V. J. Mukhatyar, A. Vakharia, S. B. Pai, B. Brahma, T. J. MacDonald, R. V. Bellamkonda, Guiding intracortical brain tumour cells to an extracortical cytotoxic hydrogel using aligned polymeric nanofibres. Nat. Mater. 13, 308–316 (2014).

10. J. G. Lyon, S. L. Carroll, N. Mokarram, R. V. Bellamkonda, Electrotaxis of Glioblastoma and Medulloblastoma Spheroidal Aggregates. Sci. Rep. (2019), doi:10.1038/s41598-019-41505-6.

11. F. K. Storm, D. L. Morton, Localized Hyperthermia in the Treatment of Cancer. CA. Cancer J. Clin. 33, 44–56 (1983).

12. I. S. Cooper, S. Stellar, Cryogenic freezing of brain tumors for excision or destruction in situ. J. Neurosurg. 20, 921–930 (1963).

13. A. A. Gage, J. M. Baust, J. G. Baust, Experimental cryosurgery investigations in vivo. Cryobiology 59, 229–243 (2009).

14. T. Fay, Early experiences with local and generalized refrigeration of the human brain. J. Neurourgery 16, 239–260 (1959).

15. M. A. Bohl, N. L. Martirosyan, Z. W. Killeen, E. Belykh, J. M. Zabramski, R. F. Spetzler, M. C. Preul, The history of therapeutic hypothermia and its use in neurosurgery. J. Neurosurg. 130, 1006–1020 (2019).

16. G. F. Rowbotham, A. L. Haigh, W. G. Leslie, Cooling cannula for use in the treatment of cerebral neoplasms. Lancet 273, 12–15 (1959).

17. D. Wion, Therapeutic dormancy to delay postsurgical glioma recurrence: the past, present and promise of focal hypothermia. J. Neurooncol. 133, 447–454 (2017).

18. X. F. Yang, B. R. Kennedy, S. G. Lomber, R. E. Schmidt, S. M. Rothman, Cooling produces minimal neuropathology in neocortex and hippocampus. Neurobiol. Dis. 23, 637–643 (2006).

19. S. G. Lomber, B. R. Payne, J. A. Horel, The cryoloop: An adaptable reversible cooling deactivation method for behavioral or electrophysiological assessment of neural function. J. Neurosci. Methods 86, 179–194 (1999).

20. H. A. Choi, N. Badjatia, S. A. Mayer, Hypothermia for acute brain injury—mechanisms and practical aspects. Nat. Rev. Neurol. 8, 214–222 (2012).

21. M. D. Smyth, S. M. Rothman, Focal Cooling Devices for the Surgical Treatment of *EpilepsyNeurosurg*. Clin. N. Am. 22, 533–546 (2011).

22. K. M. Karkar, P. A. Garcia, L. M. Bateman, M. D. Smyth, N. M. Barbaro, M. Berger, Focal Cooling Suppresses Spontaneous Epileptiform Activity without Changing the Cortical Motor Threshold. Epilepsia 43, 932–935 (2002).

23. D. Kalamida, I. V. Karagounis, A. Mitrakas, S. Kalamida, A. Giatromanolaki, M. I. Koukourakis, O. Gires, Ed. Fever-range hyperthermia vs. hypothermia effect on cancer cell viability, proliferation and HSP90 expression. PLoS One 10, e0116021 (2015).

24. Z. Matijasevic, Selective protection of non-cancer cells by hypothermia. Anticancer Res. 22, 3267–3272 (2002).

25. C. Fulbert, C. Gaude, E. Sulpice, S. Chabardès, D. Ratel, Moderate hypothermia inhibits both proliferation and migration of human glioblastoma cells. J. Neurooncol. 144, 489–499 (2019).

26. X. Zhang, Y. Lv, G. Chen, Y. Zou, C. Lin, L. Yang, P. Guo, M. Lin, Effect of mild hypothermia on breast cancer cells adhesion and migration. Biosci. Trends 6, 313–324 (2012).

27. D. K. Kelleher, C. Nauth, O. Thews, W. Krueger, P. Vaupel, Localized hypothermia: impact on oxygenation, microregional perfusion, metabolic and bioenergetic status of subcutaneous rat tumours. Br. J. Cancer 78, 56–61 (1998).

28. C. Fulbert, S. Chabardès, D. Ratel, Adjuvant therapeutic potential of moderate hypothermia for glioblastoma. J. Neurooncol. 1, 1–16 (2021).

29. L. Maggs, G. Cattaneo, A. E. Dal, A. S. Moghaddam, S. Ferrone, CAR T Cell-Based Immunotherapy for the Treatment of Glioblastoma. Front. Neurosci. 15, 535 (2021).

30. P. Hasgall, F. Di Gennaro, C. Baumgartner, E. Neufeld, B. Lloyd, M. Gosselin, D. Payne, A. Klingenböck, N. Kuster, IT’IS Database for thermal and electromagnetic parameters of biological tissues (2018), doi:10.13099/VIP21000-04-0.

31. J. R. Larkin, M. A. Simard, A. A. Khrapitchev, J. A. Meakin, T. W. Okell, M. Craig, K. J. Ray, P. Jezzard, M. A. Chappell, N. R. Sibson, Quantitative blood flow measurement in rat brain with multiphase arterial spin labelling magnetic resonance imaging. J. Cereb. Blood Flow Metab. 39, 1557–1569 (2019).

32. J. L. Boxerman, K. M. Schmainda, R. M. Weisskoff, Relative cerebral blood volume maps corrected for contrast agent extravasation significantly correlate with glioma tumor grade, whereas uncorrected maps do not. Am. J. Neuroradiol. 27, 859–867 (2006).

33. Y. Wang, L. Zhu, A. J. Rosengart, Targeted brain hypothermia induced by an interstitial cooling device in the rat neck: Experimental study and model validation. Int. J. Heat Mass Transf. 51, 5662–5670 (2008).

34. L. Zhu, C. Diao, Theoretical simulation of temperature distribution in the brain during mild hypothermia treatment for brain injury.

35. S. F. Enam, B. J. Kang, J. G. Lyon, R. V. Bellamkonda, DIY caging apparatus to facilitate chronic and continuous stimulation or recording in an awake rodent. bioRxiv, 2021.12.16.473031 (2021).

36. N. Plesnila, E. Muller, S. Guretzki, F. Ringel, F. Staub, A. Baethmann, Effect of hypothermia on the volume of rat glial cells. J. Physiol. 523 Pt 1, 155–62 (2000).

37. J. S. Bayley, C. B. Winther, K. Andersen, C. Grønkjaer, O. B. Nielsen, T. Holm Pedersen, J. Overgaard, Cold exposure causes cell death by depolarization-mediated Ca 2+ overload in a chill-susceptible insect., doi:10.1073/pnas.1813532115.

38. M. M. Salman, P. Kitchen, M. N. Woodroofe, J. E. Brown, R. M. Bill, A. C. Conner, M. T. Conner, Hypothermia increases aquaporin 4 (AQP4) plasma membrane abundance in human primary cortical astrocytes via a calcium/transient receptor potential vanilloid 4 (TRPV4)- and calmodulin-mediated mechanism. Eur. J. Neurosci. 46, 2542–2547 (2017).

39. J. Zhang, X. Xue, Y. Xu, Y. Zhang, Z. Li, H. Wang, The transcriptome responses of cardiomyocyte exposed to hypothermia. Cryobiology 72, 244–250 (2016).

40. E. C. Woolf, N. Syed, A. C. Scheck, Tumor metabolism, the ketogenic diet and ß-hydroxybutyrate: Novel approaches to adjuvant brain tumor therapy. Front. Mol. Neurosci. 9 (2016), doi:10.3389/fnmol.2016.00122.

41. T. N. Seyfried, P. Mukherjee, Targeting energy metabolism in brain cancer: Review and hypothesis Nutr. Metab. 2 (2005), doi:10.1186/1743-7075-2-30.

42. T. N. Seyfried, M. A. Kiebish, J. Marsh, L. M. Shelton, L. C. Huysentruyt, P. Mukherjee, Metabolic management of brain *cancerBiochim*. Biophys. Acta - Bioenerg. 1807, 577–594 (2011).

43. H. Sontheimer, A role for glutamate in growth and invasion of primary brain tumors. J. Neurochem. 105, 287 (2008).

44. V. F. Zhu, J. Yang, D. G. LeBrun, M. Li, Understanding the role of cytokines in Glioblastoma Multiforme pathogenesis Cancer Lett. 316, 139–150 (2012).

45. A. H. W. Nias, D. Frcr, P. M. Perry Mist, A. R. Photiou, P. Richard, Modulating the oxygen tension in tumours by hypothermia and hyperbaric oxygen. J. R. Soc. Med. 81 (1988) (available at https://www.ncbi.nlm.nih.gov/pmc/articles/PMC1291839/pdf/jrsocmed00156-0017.pdf).

46. S. Y. Lee, Temozolomide resistance in glioblastoma multiforme. Genes Dis. 3, 198–210 (2016).

47. G. Du, Y. Liu, J. Li, W. Liu, Y. Wang, H. Li, Hypothermic microenvironment plays a key role in tumor immune subversion. Int. Immunopharmacol. 17, 245–253 (2013).

48. D. Aronov, M. S. Fee, Analyzing the dynamics of brain circuits with temperature: Design and implementation of a miniature thermoelectric device. J. Neurosci. Methods 197, 32–47 (2011).

49. H. S. Venkatesh, T. B. Johung, V. Caretti, A. Noll, Y. Tang, S. Nagaraja, E. M. Gibson, C. W. Mount, J. Polepalli, S. S. Mitra, P. J. Woo, R. C. Malenka, H. Vogel, M. Bredel, P. Mallick, M. Monje, Neuronal Activity Promotes Glioma Growth through Neuroligin-3 Secretion. Cell 161, 803–816 (2015).

50. H. S. Venkatesh, W. Morishita, A. C. Geraghty, D. Silverbush, S. M. Gillespie, M. Arzt, L. T. Tam, C. Espenel, A. Ponnuswami, L. Ni, P. J. Woo, K. R. Taylor, A. Agarwal, A. Regev, D. Brang, H. Vogel, S. Hervey-Jumper, D. E. Bergles, M. L. Suvà, R. C. Malenka, M. Monje, Electrical and synaptic integration of glioma into neural circuits. Nature 573, 539–545 (2019).

51. A. J. Kalisvaart, B. Prokop, F. Colbourne, Hypothermia: Impact on plasticity following brain injury. Brain Circ. 5, 169 (2019).

52. R. F. Barth, B. Kaur, Rat brain tumor models in experimental neuro-oncology: The C6, 9L, T9, RG2, F98, BT4C, RT-2 and CNS-1 gliomas J. Neurooncol. 94, 299–312 (2009).

53. S. F. Enam, M. Calhoun, T. Saxena, S. Owen, B. Ravi, R. Chen, P. Maccarini, Devices, systems, and methods for modulating tissue temperature (2021).

54. M. R. Burns, S. Y. Chiu, B. Patel, S. G. Mitropanopoulos, J. K. Wong, A. Ramirez-Zamora, Advances and Future Directions of Neuromodulation in Neurologic Disorders., doi:10.1016/j.ncl.2020.09.004.

55. H. Wu, C. Wang, J. Liu, D. Zhou, D. Chen, Z. Liu, A. Wu, L. Yang, J. Chang, C. Luo, W. Cheng, S. Shen, Y. Bai, X. Mu, C. Li, Z. Wang, L. Chen, Evaluation of a tumor electric field treatment system in a rat model of glioma. CNS Neurosci. Ther. 26, 1168 (2020).

56. D. F. Cooke, A. B. Goldring, I. Yamayoshi, P. Tsourkas, G. H. Recanzone, A. Tiriac, T. Pan, S. I. Simon, L. Krubitzer, Fabrication of an inexpensive, implantable cooling device for reversible brain deactivation in animals ranging from rodents to primates. J. Neurophysiol. 107, 3543–3558 (2012).

57. H. Imoto, M. Fujii, J. Uchiyama, H. Fujisawa, K. Nakano, I. Kunitsugu, S. Nomura, T. Saito, M. Suzuki, Use of a Peltier chip with a newly devised local brain-cooling system for neocortical seizures in the rat. Technical note. J Neurosurg 104, 150–156 (2006).

58. H. E. Bakken, H. Kawasaki, H. Oya, J. D. W. Greenlee, M. A. Howard, A device for cooling localized regions of human cerebral cortex. J. Neurosurg. 99, 604–608 (2003).

59. S. Rockwell, In vivo-in vitro tumour cell lines: characteristics and limitations as models for human cancer. Br. J. Cancer 41, 118–122 (1980).

60. K. Riccione, C. M. Suryadevara, D. Snyder, X. Cui, J. H. Sampson, L. Sanchez-Perez, Generation of CAR T cells for adoptive therapy in the context of glioblastoma standard of care. J. Vis. Exp., e52397 (2015).

