## Supplementary Methods and Tables for "Cytostatic hypothermia and its impact on glioblastoma and survival"

### Supplementary Materials and Methods

#### a. Device fabrication

Device design and manufacturing was done in collaboration with the Pratt Bio-Medical Machine Shop at Duke University. Designs were developed on MasterCam and Fusion360 (fig. S6). Most materials and parts were obtained through McMaster, or vendors such as Digi-Key, Newark Element, Mouser, or Amazon.

*Metal and polycarbonate parts:* MasterCam designs were programmed into tool paths for a three axis CNC milling machine. Raw polycarbonate (McMaster), copper (145 copper, McMaster), and aluminum (McMaster) were used to fabricate various parts including the Interface, copper contacts, and the heat sink and its cover (fig. S6F). The tools used were .125", .0625" and 1 mm high speed steel end mills and high-speed steel drills.

*Interface:* This consisted of a polycarbonate base, a gold needle threaded through a copper part, and a thermistor with its ends wrapped around protected brass screws (fig. S6, H and I). The gold needle was fashioned from 24k 1-mm diameter gold wire (Hauser & Miller) with one end sharpened to a 45° angle by hand with a fine jewelers file in a small lathe using a 1-mm 5C collet. The gold was inserted through a hole in the adjoining copper part, ensuring the proper length for tumor penetration (3 mm from bottom surface of polycarbonate base). The gold was carefully soldered to the copper with minimal 40/60 standard solder into the top side. Care was taken to prevent (and remove) any excess solder wicking past the gold to ensure proper seating into the base recess. Remaining gold and solder on top were filed flush to the copper.
Next, two brass screws were inserted into the polycarbonate base and fixed with epoxy (Henkel Loctite, Ellsworth Adhesives). Then, under a dissection microscope, thermistors (Amphenol Advanced Sensors) that were either MRI-compatible (A96N4-GC11KA143L/37C) or MRI-incompatible (AB6N2-GC14KA143E/37C), were measured, cut, and stripped with a blade. The thermistors were threaded through a hole in the polycarbonate base on one end, and the wires were wrapped around the brass screws. Heat shrink was added around the wired screws, and then protected with a polycarbonate part and secured with additional epoxy. Prior to implantation, these were sterilized under UV light for 30 minutes and left in 70% ethanol overnight.

*Cooler:* The Cooler (fig. S6, A to E) consisted of a fan (MF20080V1-1000U-A99, Sunon), the fabricated heat sink with cover, a female connector (SM06B-GHS-TB, JST Sales America) protected by a 3D printed component (fig. S6G), a potted Peltier plate (TE-65-0.6-1.2, TeTech), and a fabricated polycarbonate plate with a copper part and steel shims (fig. S6J). The Peltier plate had thermal paste (Kryonaut, Thermal Grizzly) applied on both surfaces. For rats receiving normothermia, the Peltier plate was replaced by a block of PDMS of equivalent dimensions. An aluminum grill (9305T92, McMaster) was added above the fan to protect it from bedding. The Cooler was held together with screws.

*Wiring:* 26G wire was wrapped around screws that fastened the shims to the polycarbonate plate. These wires, along with the Peltier plate wires were passed through holes in the aluminum heatsink. The wires, including from the fan, were soldered to the female connector. The connector was protected with a 3D-printed part and then secured with epoxy and hot glue.

The device enabled the Cooler to be attached to the Interface such that the copper parts made contact to transfer heat, and the steel shims contacted the brass screws to complete the thermistor circuit (fig. S6, K and L).

**b. Animals**

All animal procedures were approved by the Duke IACUC. Fischer (CDF) and Nude (RNU) rats were purchased from Charles River at 7–9 weeks of age. All procedures began at 8–10 weeks of age.

*Tumor inoculation:* Tumor cells were grown *in vitro* for two passages, harvested the morning of the surgery, and kept on ice until injection. Animals were induced with 5% isoflurane anesthesia and maintained between 1–3% with visual monitoring of breathing. Buprenorphine-SR was administered subcutaneously at 1 mg/kg for pain control. Fur on the scalp was trimmed with an electric trimmer followed by removal with hair-removal cream. The head was then secured in a stereotaxic apparatus. Eye ointment was applied on the eyes. The scalp was sterilized with three alternating washes of 70% ethanol and chlorhexidine. Bupivicaine (0.25% w/v) was injected into the scalp. A central incision was made on the scalp and the skin retracted. A 0.6 mm conical burr was used to drill at -0.5 AP and + 3 ML to a depth of 0.8–0.9 mm. A Hamilton syringe with 26G needle loaded with at least 5 µL of either F98 cells (in Fischer rats) or U-87 MG cells (in RNU rats) was centered to the drill site. Prior to insertion, the tip was wiped of any hanging droplet. The needle was penetrated to a depth of 1.75 mm from the outer table of the skull and then retracted by 0.25 mm for a final location of 1.5 mm DV. Infusion was begun with a pump at 0.5 µL/min for 10 min. Upon 1 minute after completion, the syringe was slowly retracted, and the scalp sutured. The rat was then placed in the custom cage to begin accommodating. Supplemental nutritionally complete diet gel (76A, ClearH2O) was provided regularly.

*Implantation:* One week after inoculation, MR images were acquired was taken to confirm tumor-take. The subsequent day, the rats were prepared for implantation (fig. S8B). As previously, the rats were induced under anesthesia and buprenorphine-SR was administered subcutaneously at 1 mg/kg. The scalp was sterilized, bupivacaine (0.25% w/v) was administered, and the scalp was incised. This time, extra effort was put into retraction, scraping off the peritoneum, slightly separating the temporalis muscles, and ensuring hemostasis with etching gel (Henry Schein) and 0.4% hydrogen peroxide. Once the cranium was absent of blood and dry, burr holes were made using conical drill bits (Roboz Surgical Instruments Co.) for the gold probe, thermistor, and titanium screws (0035962, Allied Titanium). This included one burr hole with a 0.8 mm tip for the thermistor, 1.5 mm caudal from the tumor inoculation. Following this, a 1.0 mm conical burr was used 6 mm laterally from the thermistor for titanium screw (TS) 1, 4 mm caudally from TS1 for TS2, 10 mm caudally from thermistor for TS3, and 6 mm rostrally from the tumor inoculation for TS4. The original tumor inoculation burr hole was expanded to 1.4 mm. Titanium screws (filed down to be 1-mm in length) were then twisted into their holes at a depth of 0.6 mm. Next, after cleaning the skull again, the sterile Interface was gently inserted and held down while UV-curable dental cement (Henry Schein) was added to the sides and cured. Following this, layers of dental cement were added around and above the screws, Interface, and skull to secure the Interface to the skull. Upon completion, stitches were used to gently approximate the skin (including around the arms of the Interface) while keeping the surface of the Interface exposed. The rat was monitored while waking up and for 2 hours after to ensure recovery.

*Attachment:* Two days after recovery, the rat was put under anesthesia to attach the Cooler. A miniscule drop of thermal paste was added between the copper contacts and the Cooler was then screwed to the Interface. A patch cable was then connected to both the Cooler and the slipring hovering above the cage on a lever arm. For studies where MRI was possible, an additional MR image was acquired at 5–7 days after implantation (with the Cooler screwed off). After this, devices were switched on; rats with a cooling device had their fan and Peltier powered ("Device ON") while rats with normothermia only had a fan that was powered ("Device OFF"). Temperature was monitored through the Arduino connected to a computer and recorded using PuTTY v0.74 (www.putty.org).

*Monitoring and maintenance:* The intracerebral temperatures were intermittently monitored throughout the day by connecting the computer through a local network. This enabled us to respond quickly to any sudden changes in temperature, usually due to some transient failure of the patch cable, alligator clips, or device which was rectified. Over time, there was regularly an accumulation of fur inside the heat sinks and fans. This was intermittently removed with tweezers as possible. For more complicated adjustments and corrections, the rat was put under anesthesia. Rats were given nutritionally complete diet gel regularly but were also able to eat regular food and drink water from the water bottle. Supplemental treats such as softies were also provided (Bio-Serv). Cages were cleaned once weekly. Rats were typically weighed every 2–4 days but were weighed daily if weight started falling. The procedure involved transiently disconnecting the patch cable from the slipring, moving the rat to an empty cage, and subtracting the weight of the cage, patch cable, and device.

*Euthanasia:* Euthanasia criteria included: 10–15% weight loss from initial weight (after recovery from Interface implantation), or signs of significant distress, porphyrin staining around the eyes, and lack of grooming and appetite. For survival studies, rats were censored if the Interface detached from the skull. When a rat reached these criteria, they were induced under 5% anesthesia and prepared for euthanasia. The patch cable and Cooler were detached. A thoracotomy was performed followed by a trans-cardial perfusion with PBS (250 mL) and then 10% formalin (250 mL). The animals were decapitated, and the skull with the brain and Interface still in place was carefully collected and left for 24 hours in 10% formalin at 4°C. Next, the Interface and skull were carefully removed, and the brain was transferred to 20% sucrose and stored at 4°C until it sunk. Subsequently, the brain was grossly sectioned, placed on an aluminum mold and buried with Optimal Cutting Temperature compound. The aluminum portion was then exposed to liquid nitrogen to initiate snap-freezing, followed by completion on dry ice. The block was stored at -80°C until further processing.

Supplementary Table 1: Comparison of well coverage 1 wk after different durations of cytostatic hypothermia (1 to 4 wks) with 1 wk after no hypothermia (0 wks)

| Cell line | Temp (°C) | Comparison | adj. *p* -val |
|---|---|---|---|
| U-87 MG | 25 | 0 wks @D07 vs. 1 wk @D14 | 0.1811 |
| | | 0 wks @D07 vs. 2 wks @D21 | 0.0691 |
| | | 0 wks @D07 vs. 3 wks @D28 | **0.0038** |
| | | 0 wks @D07 vs. 4 wks @D35 | **<0.0001** |
| T98G | 25 | 0 wks @D07 vs. 1 wk @D14 | 0.0688 |
| | | 0 wks @D07 vs. 2 wks @D21 | **0.0005** |
| | | 0 wks @D07 vs. 3 wks @D28 | **<0.0001** |
| | | 0 wks @D07 vs. 4 wks @D35 | **<0.0001** |
| LN-229 | 25 | 0 wks @D07 vs. 1 wk @D14 | **0.0084** |
| | | 0 wks @D07 vs. 2 wks @D21 | **0.0004** |
| | | 0 wks @D07 vs. 3 wks @D28 | **0.0024** |
| | | 0 wks @D07 vs. 4 wks @D35 | **<0.0001** |
| F98 | 20 | 0 wks @D07 vs. 1 wk @D14 | **0.034** |
| | | 0 wks @D07 vs. 2 wks @D21 | 0.1849 |
| | | 0 wks @D07 vs. 3 wks @D28 | 0.1527 |
| | | 0 wks @D07 vs. 4 wks @D35 | **0.0246** |

Two-way ANOVA with Dunnet's multiple comparisons test

Supplementary Table 2: Comparison of cell circularity and size at 25°C and 20°C between Day 0 (37°C) and 3, 7, or 14 days of hypothermia

| | | | **Circularity** | | | **Average Size** | | |
|---|---|---|---|---|---|---|---|---|
| Cell line | Temp (°C) | Comparison | q | DF | adj. *p* -val | q | DF | adj. *p* -val |
| U-87 MG | 25 | D0 vs. D3 | 2.511 | 2 | 0.2378 | 7.646 | 2 | **0.032** |
| | | D0 vs. D7 | 29.13 | 2 | **0.0023** | 4.154 | 2 | 0.1012 |
| | | D0 vs. D14 | 19.2 | 2 | **0.0052** | 0.3684 | 2 | 0.962 |
| | 20 | D0 vs. D3 | 2.291 | 2 | 0.2734 | 5.96 | 2 | 0.0517 |
| | | D0 vs. D7 | 3.815 | 2 | 0.1179 | 1.231 | 2 | 0.5802 |
| | | D0 vs. D14 | 3.688 | 2 | 0.1253 | 10.64 | 2 | **0.0168** |
| T98G | 25 | D0 vs. D3 | 6.107 | 2 | **0.0494** | 6.008 | 2 | 0.0509 |
| | | D0 vs. D7 | 2.614 | 2 | 0.2234 | 2.364 | 2 | 0.2609 |
| | | D0 vs. D14 | 0.6087 | 2 | 0.8741 | 8.713 | 2 | **0.0248** |
| | 20 | D0 vs. D3 | 0.4372 | 2 | 0.9414 | 5.291 | 2 | 0.0647 |
| | | D0 vs. D7 | 2.062 | 2 | 0.3183 | 4.501 | 2 | 0.0875 |
| | | D0 vs. D14 | 1.473 | 2 | 0.485 | 0.5521 | 2 | 0.8984 |
| LN-229 | 25 | D0 vs. D3 | 238 | 2 | **<0.0001** | 6.934 | 2 | **0.0387** |
| | | D0 vs. D7 | 13.89 | 2 | **0.0099** | 15.62 | 2 | **0.0079** |
| | | D0 vs. D14 | 45.9 | 2 | **0.0009** | 2.982 | 2 | 0.1803 |
| | 20 | D0 vs. D3 | 22.67 | 2 | **0.0037** | 0.4725 | 2 | 0.9292 |
| | | D0 vs. D7 | 19.73 | 2 | **0.0049** | 0.2619 | 2 | 0.9849 |
| | | D0 vs. D14 | 25.11 | 2 | **0.0031** | 3.541 | 2 | 0.1345 |
| F98 | 25 | D0 vs. D3 | 9.847 | 2 | **0.0195** | 3.786 | 2 | 0.1195 |
| | | D0 vs. D7 | 19.96 | 2 | **0.0048** | 54 | 2 | **0.0007** |
| | | D0 vs. D14 | 25.18 | 2 | **0.003** | 22.12 | 2 | **0.0039** |
| | 20 | D0 vs. D3 | 2.563 | 2 | 0.2304 | 1.03 | 2 | 0.6712 |
| | | D0 vs. D7 | 9.258 | 2 | **0.0221** | 1.71 | 2 | 0.4078 |
| | | D0 vs. D14 | 10.58 | 2 | **0.017** | 1.037 | 2 | 0.6678 |

Two-way ANOVA with Dunnet's multiple comparisons test

Supplementary Table 3: Comparison of F98 circularity and size at Day 0 (37°C), 3, 7, or 14 days between 25°C and 20°C of hypothermia

| | | | adj. *p* -val | |
|---|---|---|---|---|
| Cell line | Day | Comparison | Circularity | Size |
| F98 | 0 | 25°C vs 20°C | 0.9985 | 0.993 |
| | 3 | 25°C vs 20°C | 0.7178 | 0.9492 |
| | 7 | 25°C vs 20°C | **0.03** | **0.0009** |
| | 14 | 25°C vs 20°C | **0.0223** | **0.028** |

Two-way ANOVA with Šídák's multiple comparisons test

Supplementary Table 4: Comparison of intracellular ATP between Day 3 at 37°C and either Day 3 or Day 7 at 25°C for each cell line

| Cell line | Comparison | adj. *p* -val |
|---|---|---|
| U-87 MG | Day 3 (37°C) vs. Day 3 (25°C) | 0.0999 |
| | Day 3 (37°C) vs. Day 7 (25°C) | **0.0122** |
| T98G | Day 3 (37°C) vs. Day 3 (25°C) | **0.0057** |
| | Day 3 (37°C) vs. Day 7 (25°C) | **0.0099** |
| LN-229 | Day 3 (37°C) vs. Day 3 (25°C) | **0.0084** |
| | Day 3 (37°C) vs. Day 7 (25°C) | **0.0006** |
| F98 | Day 3 (37°C) vs. Day 3 (25°C) | 0.1026 |
| | Day 3 (37°C) vs. Day 7 (25°C) | **0.003** |

Two-way ANOVA with Dunnet's multiple comparisons test

Supplementary Table 5: Parameters used for finite-element modelling of local intracranial hypothermia.

| **Parameter** | **Value [units]** | **Source** |
|---|---|---|
| Brain heat capacity | 3630[J/(kg*K)] | Hasgall, P. *et al*. 2018 |
| Brain density | 1046[kg/m$^3$] | Hasgall, P. *et al.* 2018 |
| Brain thermal conductivity | 0.51[W/(m*K)] | Hasgall, P. *et al.* 2018 |
| Brain blood perfusion | 0.018, 0.019333, 0.020333 [1/s] | Larkin, J. R. *et al.* 2019 |
| Tumor multiplier | 0.73, 1, 1.52, 3.96 | Boxerman, J. L. *et al.* 2006 |
| Brain/tumor metabolic heat generation | 49937[W/m$^3$] | Wang, Y. *et al.* 2008 |
| Tumor heat capacity (white brain matter) | 3583[J/(kg*K)] | Hasgall, P. *et al*. 2018 |
| Tumor density (white brain matter) | 1041[kg/m$^3$] | Hasgall, P. *et al.* 2018 |
| Tumor thermal conductivity (white brain matter) | 0.48[W/(m*K)] | Hasgall, P. *et al.* 2018 |

**Supplementary Table 6: Histological analysis of immediate region adjacent to probe.** Brain sections stained with hematoxylin and eosin were analyzed by an expert neuropathologist who was blinded to the groups. After analysis, the data were organized based on treatment groups and labeled "Hypo" for rats that had their devices switched on and "Normo" for rats who had their devices switched off.

| Treatment | ID | Periprobe | Areas away from probe not contiguous with periprobe reaction |
|---|---|---|---|
| Hypo | F03 | Necrosis with inflammation | no definitive tumor away from probe reaction; deeper tumor shows focal tumor-intrinsic necrosis |
| Hypo | F05 | Necrosis with inflammation | no definitive tumor away from probe reaction; deeper tumor shows extensive tumor-intrinsic necrosis |
| Hypo | F07 | Necrosis with inflammation | no definitive tumor away from probe reaction on sections provided |
| Hypo | F08 | Necrosis with inflammation | Small foci of tumor-intrinsic necrosis; deeper tumor shows extensive tumor-intrinsic necrosis |
| Hypo | F12 | Necrosis with inflammation | no definitive tumor away from probe reaction; deeper tumor shows extensive tumor-intrinsic necrosis |
| Hypo | F13 | Necrosis with inflammation | rim of viable tumor |
| Normo | F01 | probe cavity centered in tumor-intrinsic necrosis | probe cavity centered in large area of tumor-intrinsic necrosis |
| Normo | F04 | Minimal inflammation | large areas of tumor-intrinsic necrosis |
| Normo | F06 | probe cavity centered in tumor-intrinsic necrosis | probe cavity centered in large area of tumor-intrinsic necrosis |
| Normo | F09 | Minimal inflammation | large areas of tumor-intrinsic necrosis |
| Normo | F10 | probe cavity not intact on sections | large areas of tumor-intrinsic necrosis |
| Hypo | N04 | Necrosis with inflammation | no definitive tumor away from probe reaction on sections provided |
| Hypo | N05 | Necrosis with inflammation | no definitive tumor away from probe reaction; deeper tumor shows focal necrosis and large cystic areas |
| Hypo | N06 | Necrosis with inflammation | viable tumor |
| Hypo | N07 | Necrosis with inflammation | no definitive tumor away from probe reaction on sections provided |
| Hypo | N09 | Necrosis with inflammation | viable tumor |
| Hypo | N10 | Necrosis with inflammation | no definitive tumor away from probe reaction on sections provided |
| Normo | N02 | Minimal inflammation; focal mineralized material (?skull) | Small foci of tumor-intrinsic necrosis |
| Normo | N03 | Minimal inflammation | Small foci of tumor-intrinsic necrosis |
| Normo | N08 | Minimal inflammation | large areas of tumor-intrinsic necrosis |
| Normo | N11 | Minimal inflammation | large areas of tumor-intrinsic necrosis |
| Normo | N12 | Minimal inflammation | large areas of tumor-intrinsic necrosis |

**General observations:**

"Small foci of tumor-intrinsic necrosis" show pyknosis (apotosis?) (early ischemic?); associated with at least poly inflammation (can't distinguish apoptosis from mono inflammatory) cells

"Large areas of tumor-intrinsic necrosis" are sometimes geographic/peritheliomatous/cystic, with at least poly inflammation

Periprobe necrosis includes abundant neutrophils around probe cavity, and macrophages more evident further away

"no definitive tumor away from probe reaction on sections provided" means cannot distinguish tumor cells from reactive cells; it does not mean no tumor

The dense DAPI populations around the probe cavity are inflammatory cells; mostly polys, sometimes CD45+ (these may be macrophages or lymphocytes)

**Supplementary Table 7: Histological analysis of peritumoral and intratumoral regions.** Brain sections stained with either hematoxylin and eosin (H&E) or immunohistochemical markers were analyzed by a Neuropathologist who was blinded to the treatment groups. After analysis, the data were organized based on treatment groups and labeled "Hypo" for rats that had their devices switched on and "Normo" for rats who had their

| | | Peritumoral findings on H&E | | | | Intratumoral findings on H&E | | | Peritumoral (PT) and neuropil (NP) immunofluorescent staining | | | | |
|---|---|---|---|---|---|---|---|---|---|---|---|---|---|
| **Treatment** | **ID** | **edema** | **hemorrhage** | **hypercellularity** | **vascular changes** | **necrosis** | **VRSi** | **tumor size** | **GFAP gliosis** | **Ki67** | **NeuN** | **CD45** | **CC3** |
| Hypo | F03 | moderate | _ | moderate | moderate | yes | yes | medium | diffuse to edge of cortex | CTT | intact | mod PT, minimal NP | minimal NP |
| Hypo | F05 | moderate | _ | mild | moderate | yes | yes | medium | extensive around tumor | CTT | intact | mild PT, minimal NP | _ |
| Hypo | F07 | moderate | _ | moderate | moderate | yte | _ | small | extensive around tumor | scatter @ border | intact | mild PT | mild NP |
| Hypo | F08 | moderate | _ | mild | moderate | _ | yes | large | extensive around tumor | few @ border | intact | mild PT | stain failed |
| Hypo | F12 | marked | _ | moderate | moderate | yes | yes | medium | peritumoral | few @ border | intact | mod PT, minimal NP | mild NP |
| Hypo | F13 | moderate | _ | moderate | moderate | yes | _ | small | peritumoral | scatter @ border | intact | mod PT, minimal NP | mild NP, and PT |
| Normo | F01 | moderate | _ | mild | _ | yes | yes | large | diffuse to edge of cortex | CTT | intact | dense PT, minimal NP | minimal NP |
| Normo | F04 | moderate | _ | mild | _ | yes | _ | large | extensive around tumor | CTT | intact | dense PT, minimal NP | _ |
| Normo | F06 | moderate | _ | moderate | _ | yes | yes | medium | diffuse to edge of cortex | CTT | intact | mod PT, mild NP | _ |
| Normo | F09 | moderate | mild fresh | marked | _ | yes | yes | medium | extensive around tumor | few @ border | intact | mod PT, minimal NP | mild NP |
| Normo | F10 | moderate | _ | moderate | mild | yes | yes | medium | stain failed | stain failed | stain failed | stain failed | stain failed |
| Hypo | N04 | mild | _ | moderate | mild | yes | _ | small | peritumoral | scatter @ border | intact | stain failed | mild NP, and PT |
| Hypo | N05 | moderate | _ | mild | mild | yes | _ | small | extensive around tumor | scatter @ border | intact | stain failed | _ |
| Hypo | N06 | mild | mild fresh | mild | _ | yes | _ | small | peritumoral | CTT | intact | stain failed | minimal NP |
| Hypo | N07 | _ | mild fresh | _ | mild | yes | _ | small | diffuse to edge of cortex | scatter @ border | intact | stain failed | minimal NP |
| Hypo | N09 | mild | focal old | mild | moderate | yes | _ | medium | peritumoral | CTT | intact | stain failed | minimal NP |
| Hypo | N10 | mild | _ | mild | mild | yes | _ | small | peritumoral | CTT | intact | mild PT | minimal NP |
| Normo | N02 | mild | _ | mild | mild | _ | _ | large | peritumoral | stain failed | intact | stain failed | _ |
| Normo | N03 | _ | _ | _ | _ | _ | _ | large | peritumoral | CTT | intact | minimal PT | minimal NP |
| Normo | N08 | _ | _ | _ | _ | yes | _ | large | diffuse to edge of cortex | CTT | intact | stain failed | minimal NP |
| Normo | N11 | mild | _ | mild | _ | yes | _ | large | peritumoral | CTT | intact | minimal PT | minimal NP |
| Normo | N12 | _ | _ | _ | _ | yes | _ | large | peritumoral | stain failed | intact | stain failed | minimal NP |

**General observations:**
No evidence of peritumoral or distant infarction or vascular thrombosis in any brain
No evidence of infection (no neutrophilic encephalitis or meningitis, no bacterial or fungal colonies evident on H&E) in any brain
No evidence of herniation (no effacement of CSF spaces) in any brain
No loss of NeuN staining; intact to border of tumor on all sections
No evidence of diffuse infiltration of glioma
Vascular changes: microvessel dilation and/or hyperplasia
All reactions are ipsilateal to tumor, including reactive gliosis; no contralateral effect other than mass effect
Ki67 is largely confined to tumor (CTT) except where otherwise noted
VRSi = Virchow-Robin space involvement
CD45 will stain macrophages, lymphocytes and microglia
From one area to another, features can change a level, from none to mild to moderate to marked, so sampling error must be considered
